# Focused framework sampling recovers binding-positive humanized anti-amyloid-β antibodies in a single sorting round

**DOI:** 10.64898/2026.08.14.744767

**Authors:** Youngju Kim, Hayoung Kwon, Jimin Song, Yujin Lee, Minho Park, Chang-Han Lee

## Abstract

Therapeutic antibody development requires workflows that integrate antigen-reactive clone discovery with efficient humanization and early developability assessment. Here, we combined immune yeast fragment antigen-binding (Fab) display with single-round focused humanization and applied the workflow to antibodies against amyloid-β (Aβ)-derived preparations. Immunization with Aβ_1-42_ aggregate preparations generated a Fab-display library with a diversity of approximately 3.5 × 10^8^. Magnetic enrichment followed by fluorescence-activated cell sorting (FACS) identified three sequence-distinct immunoglobulin G (IgG)-format candidates, of which CLAB17 and CLAB45 were advanced to humanization. Structure-guided libraries sampled framework positions predicted to support complementarity-determining regions (CDRs) or heavy-and light-chain variable-domain packing, and a single FACS round recovered binding-positive variants CLAB17-h2 and CLAB45-h8. Both retained the parental CDRs and showed increased predicted humanness, favorable computational developability triage profiles, and high purity by sodium dodecyl sulfate-polyacrylamide gel electrophoresis (SDS-PAGE). By enzyme-linked immunosorbent assay (ELISA), CLAB17-h2 showed a lower apparent half-maximal effective concentration (EC_50_) for Aβ_1-42_AggreSure, whereas CLAB45-h8 showed a lower apparent EC_50_ for pyroglutamate-modified Aβ_3-42_ (Aβ_pE3-42_). Because the preparations were not resolved into defined assembly states, these antibodies are considered Aβ-preparation-binding rather than aggregate-state-selective candidates. This workflow provides a practical route from immune-repertoire discovery to binding-positive humanized antibodies.

## Introduction

Alzheimer’s disease (AD) is characterized by the accumulation of amyloid-β (Aβ), which remains a major therapeutic target in AD (Hampel et al., 2021; Hardy and Selkoe, 2002; Karran and De Strooper, 2016). Aβ exists as a heterogeneous continuum of conformational and assembly states, ranging from monomers and soluble oligomers to protofibrils and fibrils, with substantial differences in structure and surface-exposed epitopes (Benilova et al., 2012; Ono and Watanabe-Nakayama, 2021; Selkoe, 2008). The aggregation-prone Aβ_1-42_ species is particularly relevant to this heterogeneity and has been widely used as a target for antibody discovery (Hur, 2022; Ono and Watanabe-Nakayama, 2021). Consequently, the molecular form of Aβ presented during antibody discovery can strongly influence the reactivity profile of recovered antibodies.

Building on preclinical work showing that peripherally administered anti-Aβ antibodies reduce amyloid pathology, neutralize synaptotoxic oligomers, and promote antibody-mediated clearance (Bard et al., 2000; Illouz et al., 2021; Klyubin et al., 2005), clinical studies of anti-Aβ monoclonal antibodies further illustrate the importance of Aβ species reactivity. Lecanemab and donanemab reduce cerebral amyloid burden and modestly slow cognitive and functional decline in early AD (van Dyck et al., 2023; Sims et al., 2023), yet they engage different Aβ forms. Lecanemab preferentially recognizes soluble protofibrillar Aβ, whereas donanemab targets pyroglutamate-modified Aβ enriched in deposited plaques (DeMattos et al., 2012; Shukla and Misra, 2024; Söderberg et al., 2023). Differences in Aβ-form engagement have also been discussed in relation to amyloid-related imaging abnormalities (ARIA), the principal dose-limiting adverse event of this antibody class (Hampel et al., 2021; Yaari et al., 2022). These precedents highlight the value of antibody-discovery strategies that can generate and characterize binders against structurally heterogeneous Aβ-derived materials.

A practical challenge, however, is to connect antibody discovery efficiently with subsequent humanization. Yeast surface display enables flow-cytometry-based selection in which antibody display and antigen binding can be monitored simultaneously, allowing controlled enrichment of antigen-reactive clones (Chao et al., 2006; Doerner et al., 2014). Immune display libraries additionally capture antibodies that have undergone in vivo repertoire selection and affinity maturation, while retaining the throughput and experimental control of in vitro screening (Schröter et al., 2018). Antibodies recovered from murine immune repertoires nevertheless require humanization for therapeutic development. Conventional complementarity-determining region (CDR) grafting can compromise antigen binding because framework residues, including residues within the vernier zone and the heavy-chain variable (VH) and light-chain variable (VL) packing interface, can influence CDR conformation and antigen-binding-site geometry (Foote and Winter, 1992; Shembekar et al., 2014). Humanization therefore represents a distinct engineering bottleneck after antibody discovery, particularly when a fixed CDR-grafted framework does not preserve the structural context required for binding. Focused experimental sampling of selected framework residues provides a potential means to identify humanized framework combinations that retain the parental CDRs and antigen reactivity without requiring extensive iterative redesign.

In this study, we established an integrated immune yeast Fab-display and focused humanization workflow using Aβ_1-42_-derived preparations as a model antigen system. An immune Fab-display library generated from Aβ_1-42_-immunized mice was enriched by sequential magnetic-activated cell sorting (MACS) and fluorescence-activated cell sorting (FACS), followed by immunoglobulin G (IgG)-format validation of selected antibodies. Two sequence-distinct antibody lineages were subsequently subjected to structure-guided focused humanization in which selected framework positions were experimentally sampled while preserving the parental CDRs. Single-round FACS screening recovered humanized variants that retained apparent Aβ-preparation binding and showed increased predicted humanness and favorable computational developability triage profiles. Together, this study provides a practical workflow linking immune-repertoire antibody discovery with rapid, single-round humanization.

## Materials and Methods

Detailed protocols, reagents, and catalogue numbers are provided in the Supplementary Methods.

### Aβ_1-42_-derived antigens and mouse immunization

Aβ_1-42_aggregate preparations were generated from hexafluoroisopropanol (HFIP)-treated peptide as described (Stine et al., 2011), by incubation at 100 μM in phosphate-buffered saline (PBS, pH 7.4) at 37 °C for 7 days. Preparations were not fractionated into defined monomeric, oligomeric, protofibrillar, or fibrillar species, so binding data are reported as reactivity to the tested Aβ-derived preparations. Immunization and serum enzyme-linked immunosorbent assay (ELISA) followed established laboratory protocols (Chung et al., 2026; Kang et al., 2024; Lim et al., 2026). Five six-week-old BALB/c mice received 50 μg of preparation in alhydrogel (1:1, v/v) intraperitoneally at weeks 1, 3, 5, 7, and 9; spleen and bone marrow were collected after the final boost. Animal procedures were approved by the Institutional Animal Care and Use Committee of Seoul National University (SNU-230519-9-3).

### Immune yeast Fab-display library construction and screening

Murine VH and VL repertoires amplified from spleen and bone marrow complementary DNA (cDNA) (**Supplementary Table S1**) were fused to the first heavy-chain constant domain (CH1) and the light-chain constant region (CL), respectively, and displayed as diploid Fab libraries on the mating-based pYDS-H/JAR200 and pYDS-K/YvH10 yeast platform described previously (Kim et al., 2026); library diversity and mating efficiency were measured by differential selective plating (**Fig. 1D**). One MACS round (1 × 10^9^ induced cells, 100 nM biotinylated Aβ_1-42_ aggregate probe) was followed by a single FACS round (50 nM) collecting the top 1% antigen-binding fraction of Fab-displaying cells. Enrichment was monitored with biotinylated Aβ_1-42_ aggregate and Aβ_1-42_ AggreSure probes across the pre-sort (R1), MACS-enriched (R2), and FACS-sorted (R3) populations (**Fig. 2A**). Single clones from R3 were screened at 30 nM of each probe and sequenced by colony polymerase chain reaction (PCR).

**Fig. 1.**
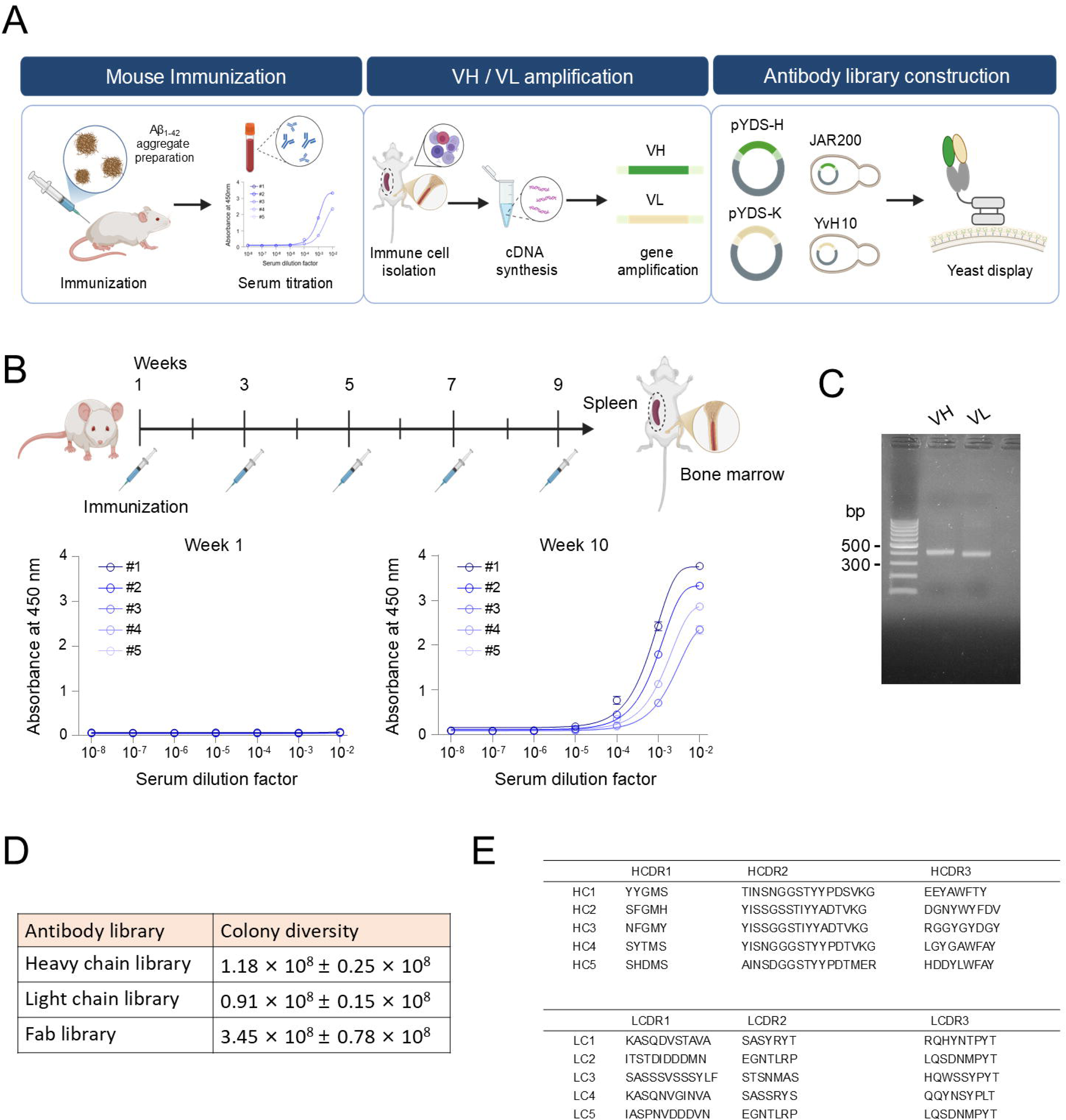
Immunization with amyloid-β_1-42_ (Aβ_1-42_) aggregate preparations and construction of an immune yeast fragment antigen-binding (Fab)-display library. **(A)** Schematic of the antibody-discovery workflow. Mice were immunized with Aβ_1-42_ aggregate preparations; spleen and bone marrow were harvested for immune-cell isolation, complementary DNA (cDNA) synthesis, heavy-chain variable/light-chain variable (VH/VL) amplification, and yeast Fab-display library construction. **(B)** Immunization schedule and serum enzyme-linked immunosorbent assay (ELISA). Sera from five mice were serially diluted and tested against immobilized Aβ_1-42_ aggregate preparation; serum dilution factor (x-axis) versus absorbance at 450 nm (y-axis). **(C)** Representative polymerase chain reaction (PCR) amplification of murine VH and VL repertoires from immune-derived cDNA; molecular-weight markers are shown in base pairs. **(D)** Colony diversity of the heavy-chain, light-chain, and paired Fab libraries. Mean ± standard deviation (SD). **(E)** Representative complementarity-determining region (CDR) sequences sampled after library construction, illustrating recovery of multiple VH/VL CDR compositions; not used to quantify full repertoire diversity.

**Fig. 2.**
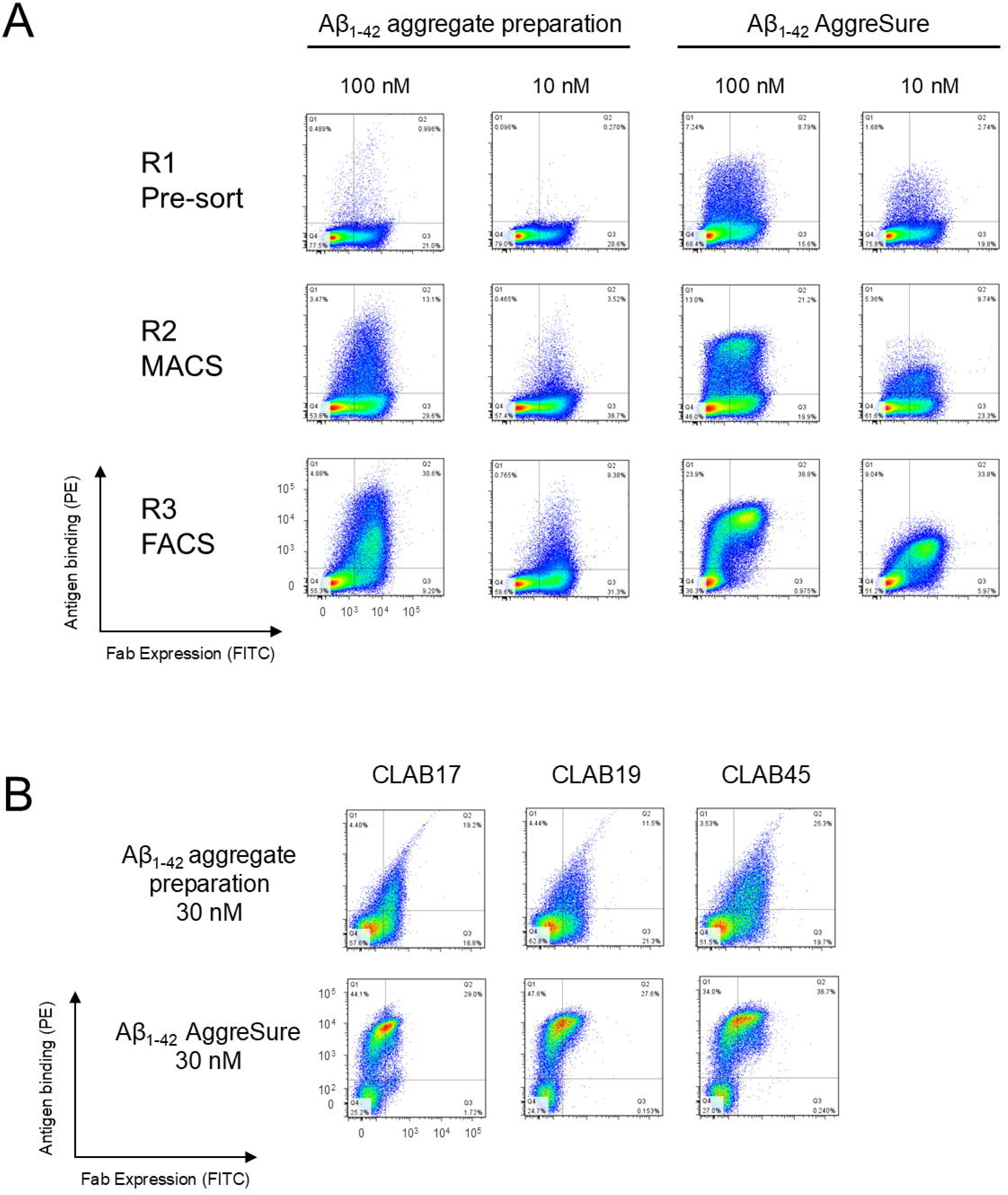
Yeast fragment antigen-binding (Fab)-display screening enriches clones binding amyloid-β (Aβ)-derived screening probes. **(A)** Flow-cytometric analysis of library enrichment using biotin-Aβ_1-42_aggregate probe and in-house-biotinylated Aβ_1-42_AggreSure probe at the indicated concentrations. Fab expression, fluorescein isothiocyanate (FITC); antigen binding, phycoerythrin (PE). R1, pre-sort library; R2, magnetic-activated cell sorting (MACS)-enriched; R3, after one fluorescence-activated cell sorting (FACS) round. Values reported in the text correspond to the displayed Q2 FITC-positive/PE-positive fractions among gated singlets. Q2 fractions increased from 0.996% to 13.1% and 30.6% for Aβ_1-42_aggregate at 100 nM, from 0.270% to 3.52% and 9.38% for Aβ_1-42_ aggregate at 10 nM, from 8.79% to 21.2% and 38.8% for Aβ_1-42_AggreSure at 100 nM, and from 2.74% to 9.74% and 33.8% for Aβ_1-42_ AggreSure at 10 nM across R1, R2, and R3, respectively. **(B)** Representative single-clone FACS of selected R3-derived clones stained with Aβ_1-42_ aggregate preparation and Aβ_1-42_ AggreSure preparation at 30 nM. CLAB17, CLAB19, and CLAB45 were prioritized for immunoglobulin G (IgG)-format validation based on combined Fab display, antigen-binding signal, staining reproducibility, sequence distinction, and IgG-conversion feasibility.

### IgG expression and ELISA-based binding analysis

Selected VH/VL pairs were reformatted as chimeric or humanized human IgG1/κ, expressed in human embryonic kidney 293F (HEK293F) cells, and purified on Protein A (Lim et al., 2026). Binding to immobilized Aβ_1-42_ AggreSure and pyroglutamate-modified Aβ_3-42_ (Aβ_pE3-42_) was measured by ELISA as described (Yang et al., 2026), with horseradish peroxidase (HRP)-conjugated anti-human IgG fragment crystallizable (Fc) detection. Apparent half-maximal effective concentration (EC_50_) values are assay-dependent, avidity-influenced readouts, not monovalent affinity constants. Assays with the wild-type Fc (wtFc)-Aβ_1-42_ fusion probe and non-target proteins are described in the Supplementary Methods.

### Design and screening of focused humanization libraries

Human acceptor frameworks were selected as the human germline V genes with the highest sequence identity to the parental murine V regions, and the parental CDRs of CLAB17 and CLAB45 were grafted onto them. Variable fragment (Fv) models of the parental murine and CDR-grafted antibodies were generated with AlphaFold 3 (Abramson et al., 2024) and superposed (**Fig. 4B,C**). Framework positions predicted to influence CDR support or VH/VL packing were then sampled between the human and the corresponding parental murine residue, with all parental CDRs preserved (**Fig. 4D**; **Table 2**). Libraries were constructed and mated as above and sorted in a single FACS round using Aβ_1-42_ AggreSure at 100 and 10 nM. Ten clones per lineage were screened at 30 nM, and final VH/VL sequences were determined (**Table 2**).

**Fig. 3.**
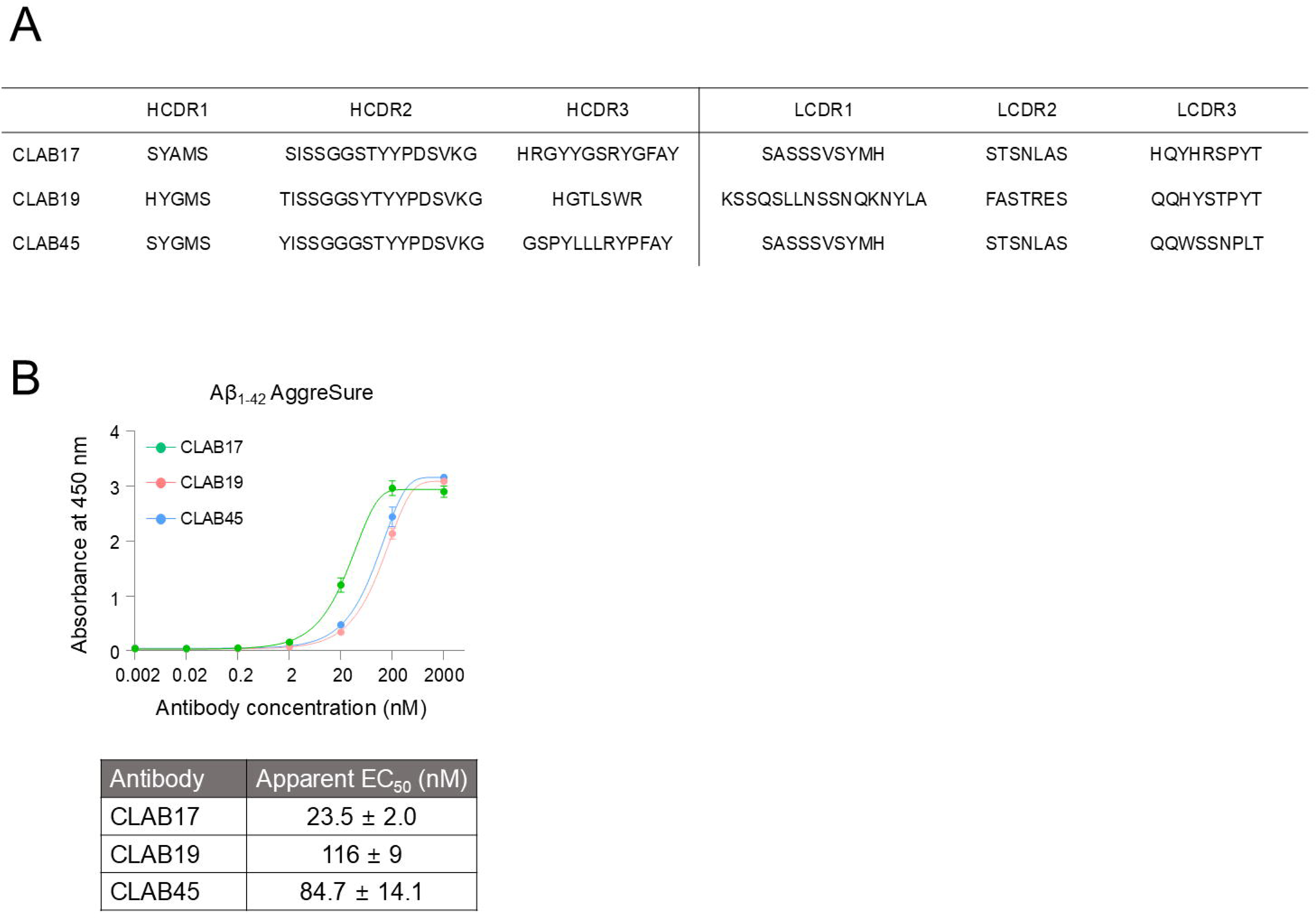
Immunoglobulin G (IgG)-formatted CLAB candidates show apparent enzyme-linked immunosorbent assay (ELISA) binding to immobilized amyloid-β_1-42_ (Aβ_1-42_) AggreSure preparation. **(A)** Complementarity-determining region (CDR) sequences of CLAB17, CLAB19, and CLAB45, differing particularly in heavy-chain CDR3 (HCDR3) and light-chain CDR3 (LCDR3), indicating non-identical lineages. **(B)** Binding of IgG-formatted CLAB17, CLAB19, and CLAB45 to immobilized Aβ_1-42_ AggreSure preparation. Apparent half-maximal effective concentration (EC_50_) values were calculated from dose-response curves and are reported as apparent ELISA EC_50_ values, not monovalent affinity constants. Data are shown as mean ± standard deviation (SD).

**Fig. 4.**
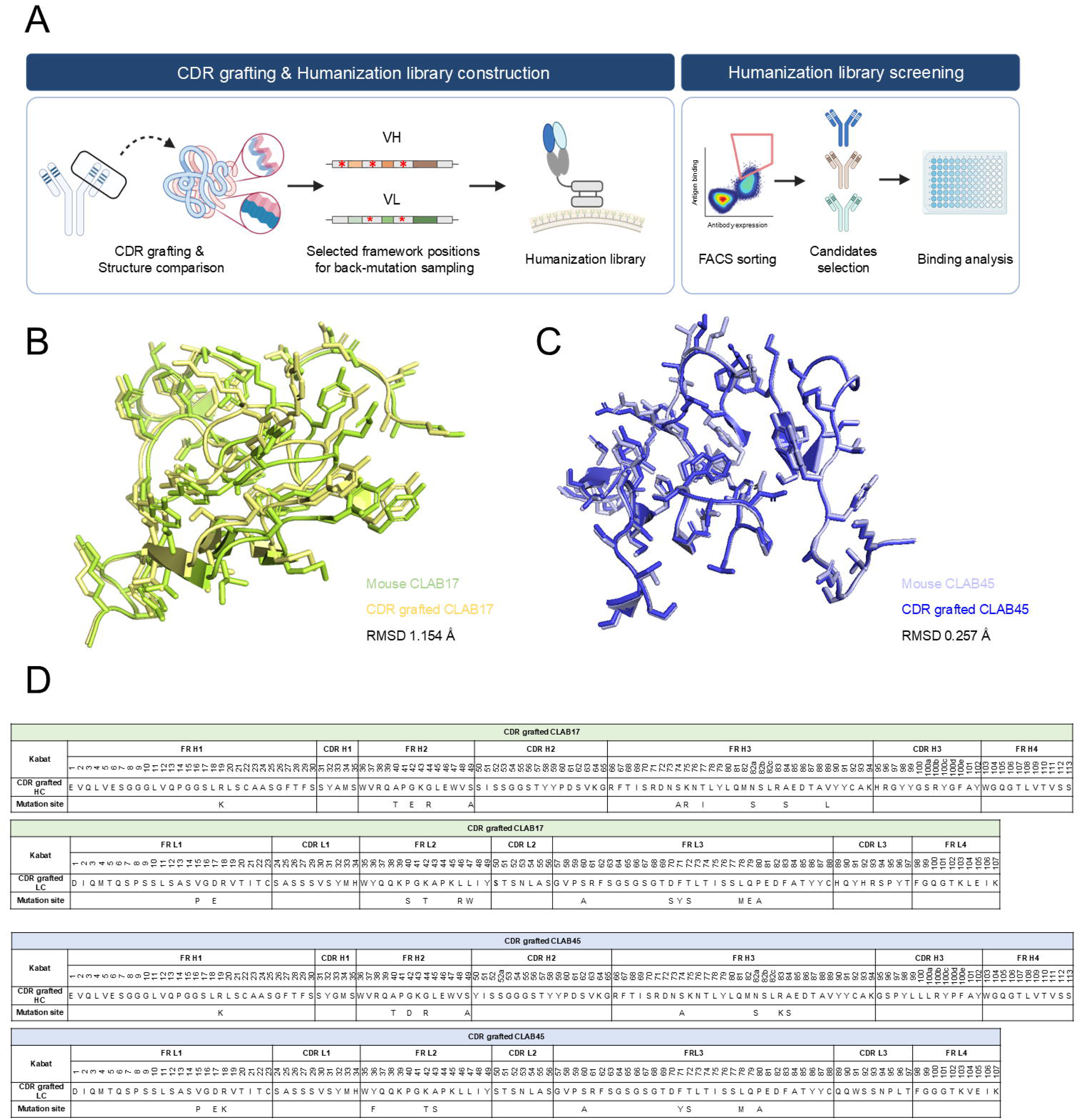
Structure-guided design of focused humanization libraries for CLAB17 and CLAB45. **(A)** Humanization strategy: complementarity-determining region (CDR) grafting onto human templates, structural comparison, focused framework back-mutation design at selected framework positions, single-round fluorescence-activated cell sorting (FACS) library screening, candidate selection, and binding analysis. (B,C) Structural overlay of parental murine and CDR-grafted variable fragment (Fv) models for CLAB17 **(B)** and CLAB45 **(C)**. Whole-Fv backbone root-mean-square deviation (RMSD) was 1.154 Å for CLAB17 and 0.257 Å for CLAB45. These values were used qualitatively to guide focused humanization-library design. **(D)** Sequence maps of CDR-grafted CLAB17 and CLAB45 variants showing framework regions, CDRs, Kabat numbering, and selected framework positions included in the focused humanization-library design. The “Mutation site” row indicates parental murine framework residues included as back-mutation options during library construction. Final selected sequences are shown in Fig. 5C, and position-resolved residue retention is summarized in **Table 2**.

**Table 1.**
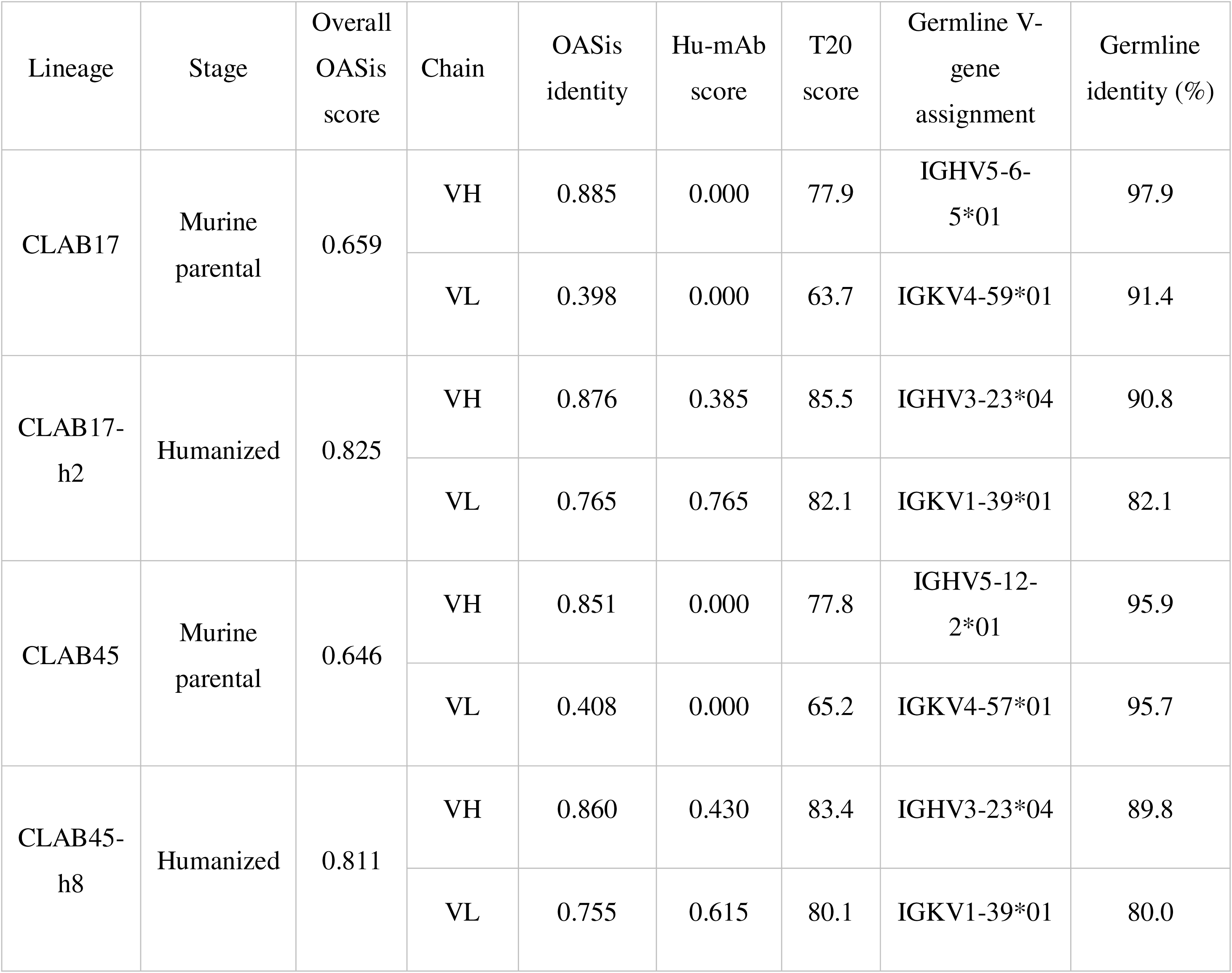
Predicted humanness scores and species-appropriate germline assignments of parental murine and final humanized CLAB antibodies. OASis overall score, chain-specific OASis identity, Hu-mAb score, T20 score, germline V-gene assignment, and germline identity are shown for parental murine antibodies and the final selected humanized variants. Germline assignments were determined using species-appropriate international ImMunoGeneTics information system (IMGT) germline databases: parental murine antibodies were assigned against mouse germline genes, and final humanized variants were assigned against human germline genes. Complementarity-determining region (CDR)-grafted sequences were used as design intermediates for focused humanization-library construction and were not included in this final comparison. OASis, Observed Antibody Space identity search; Hu-mAb, humanness scoring and humanization method; T20, antibody humanness score; VH, heavy-chain variable domain; VL, light-chain variable domain.

**Table 2.**
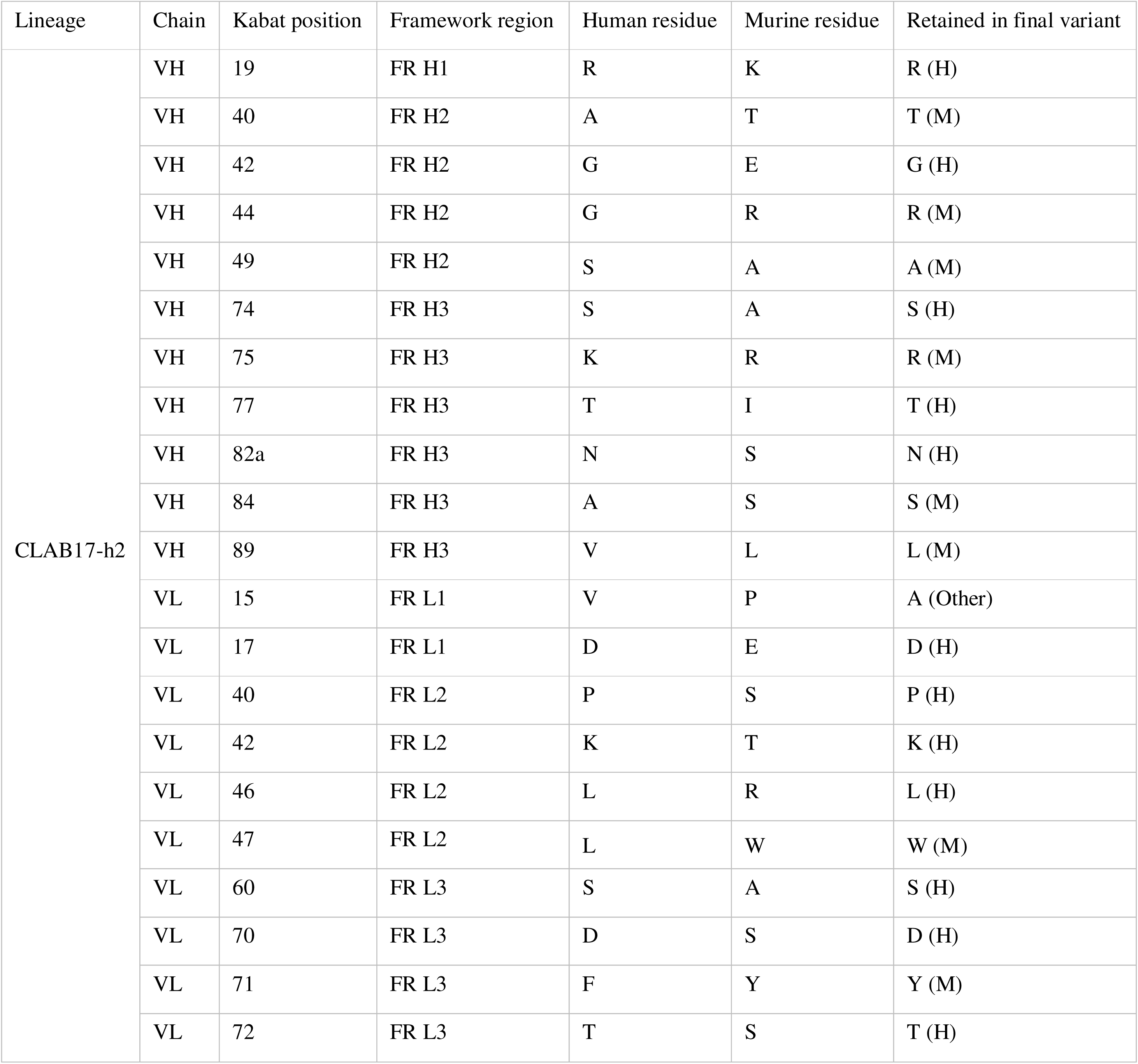
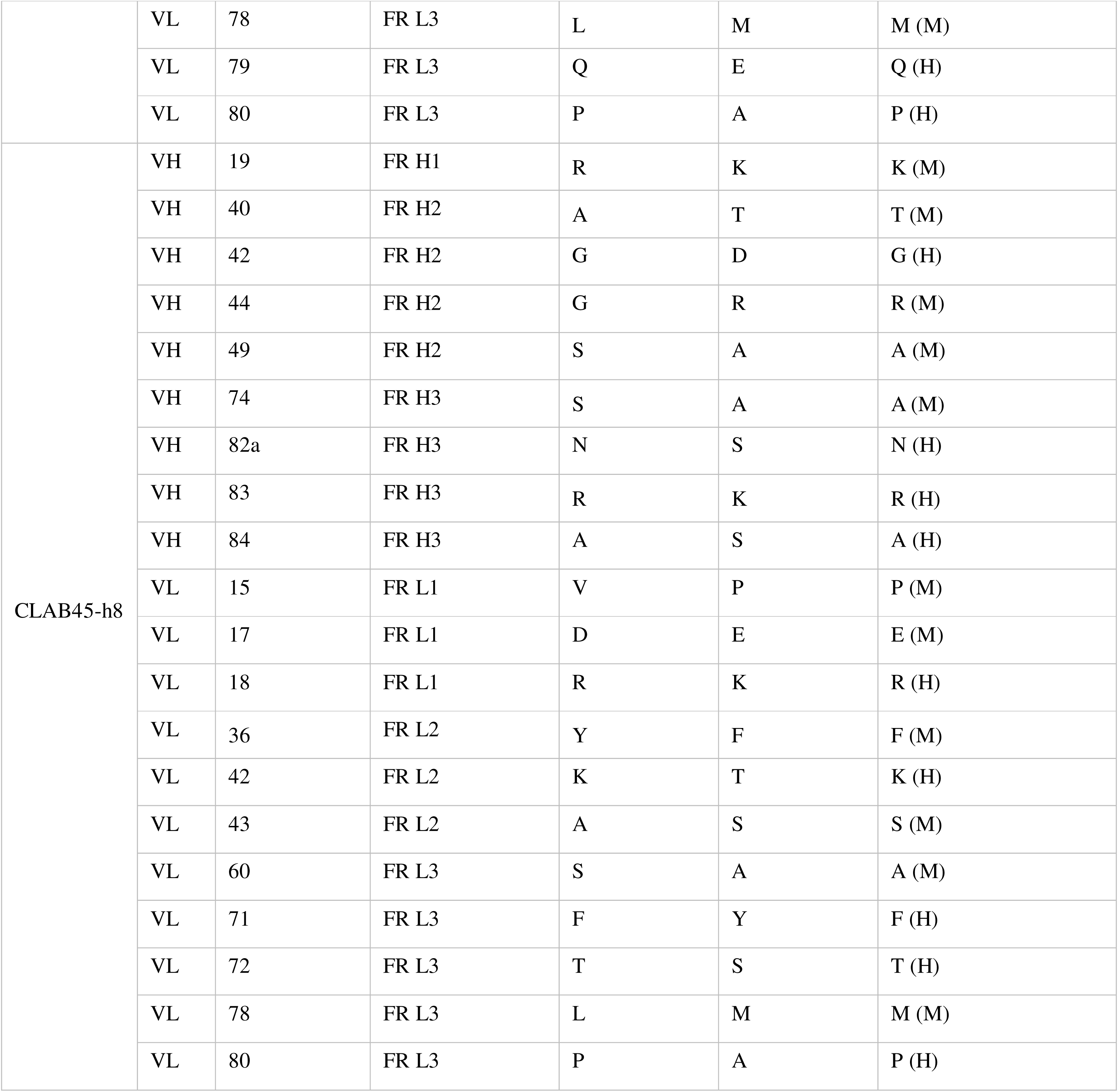
Selected framework positions sampled in the focused humanization libraries and residues retained in the final humanized variants. For each sampled framework position, the human framework residue in the complementarity-determining region (CDR)-grafted template, the parental murine residue included as a back-mutation option, and the residue identified in the final sequenced variant are listed. The sampled positions represent a focused subset of framework residues selected from regions predicted to influence CDR support or heavy-chain variable/light-chain variable (VH/VL) domain packing; they are not intended to constitute an exhaustive annotation of all vernier-zone or packing residues. H indicates retention of the human framework residue, M indicates retention of the parental murine residue, and Other indicates a residue distinct from both the human framework and parental murine residues. Residues in the final-variant column were assigned directly from the sequenced CLAB17-h2 and CLAB45-h8 VH/VL amino-acid sequences after Kabat numbering and comparison with the CDR-grafted human and parental murine frameworks; unannotated positions in the sequence maps were not assumed to be human. CLAB17-h2 VL Kabat position 15 retained alanine, which was distinct from both the CDR-grafted human residue valine and the parental murine residue proline, and was therefore annotated as Other.

### Humanness, germline, and developability analyses

Humanness was scored with BioPhi OASis (Observed Antibody Space identity search) (Prihoda et al., 2022), Hu-mAb (Marks et al., 2021), and T20 (Gao et al., 2013), and germlines were assigned by Immunoglobulin BLAST (IgBLAST) (Ye et al., 2013) against species-appropriate international ImMunoGeneTics information system (IMGT) databases (Lefranc et al., 2015). Final sequences were scanned in silico for common sequence liabilities and evaluated with the Therapeutic Antibody Profiler (TAP) for total CDR length, CDR-vicinity patches of surface hydrophobicity (PSH), positive charge (PPC), and negative charge (PNC), and the structural Fv charge symmetry parameter (SFvCSP) (Raybould et al., 2019).

## Data analysis

Data are mean ± standard deviation (SD); no significance testing was performed. Apparent EC_50_ values were calculated with the AAT Bioquest EC_50_ calculator, graphs prepared in GraphPad Prism 9, and schematics created with BioRender.com.

## Results

### Construction of an immune yeast Fab library from mice immunized with Aβ_1-42_ aggregate preparations

To isolate antibodies binding the Aβ preparations used in this study, mice were immunized with Aβ_1-42_ aggregate preparations and monitored for serum reactivity by ELISA (**Fig. 1A**). Hereafter, “Aβ_1-42_ aggregate preparation” denotes the immunogen and library-generation material, and “Aβ_1-42_ AggreSure preparation” the principal screening and validation probe.

Sera from five immunized mice were analyzed over dilutions from 10^−2^ to 10^−8^. Serum reactivity against the Aβ_1-42_ aggregate preparation was low at baseline and increased after the immunization course, indicating an Aβ-preparation-reactive humoral response suitable for repertoire recovery (**Fig. 1B**). Spleen and bone marrow were harvested as repertoire sources; messenger RNA-derived cDNA was used to amplify murine VH and VL repertoires (**Fig. 1C**). The heavy-chain library reached a colony diversity of 1.18 × 10^8^ ± 0.25 × 10^8^ and the light-chain library 0.91 × 10^8^ ± 0.15 × 10^8^. After pairing by yeast mating (mating efficiency 51.8%), the Fab library reached 3.45 × 10^8^ ± 0.78 × 10^8^ (**Fig. 1D**). Representative VH/VL sequencing confirmed recovery of multiple distinct CDR compositions across heavy-chain CDRs (HCDR1–3) and light-chain CDRs (LCDR1–3) (**Fig. 1E**), establishing an immune-derived Fab library for screening against Aβ-preparation probes.

### Yeast Fab-display screening enriched clones binding the Aβ preparations used as screening probes

Selection was performed with the biotinylated Aβ_1-42_ aggregate probe, and enrichment was monitored with both the aggregate and Aβ_1-42_ AggreSure probes, with Fab expression and antigen binding measured simultaneously by flow cytometry (**Fig. 2A**). The double-positive fraction in the upper-right quadrant (Q2), corresponding to cells positive for both Fab display and antigen binding, increased progressively across the pre-sort (R1), MACS-enriched (R2), and FACS-sorted (R3) populations. At 100 nM, Q2 rose from 0.996% to 30.6% for the biotinylated Aβ_1-42_ aggregate probe and from 8.79% to 38.8% for the Aβ_1-42_ AggreSure probe, with the same trend at 10 nM. MACS enrichment followed by one FACS round therefore increased the representation of double-positive Aβ-preparation-binding Fab-displaying yeast cells across both analytical probes and concentrations.

From R3, 49 single clones were isolated and analyzed by single-clone FACS on two parameters, Fab expression and antigen binding (**Fig. 2B** and **Supplementary Fig. S1**). Clones were prioritized for IgG conversion based on Fab display, antigen-binding signal, staining reproducibility, sequence distinction, and IgG-conversion feasibility. After IgG-format validation, CLAB17 and CLAB45 were advanced as principal candidates because they showed stronger apparent ELISA binding and represented distinct CDR architectures, whereas CLAB19 was retained as a weaker comparator and was not humanized.

### IgG-formatted CLAB candidates show apparent ELISA binding to immobilized Aβ_1-42_ AggreSure preparation

CLAB17, CLAB19, and CLAB45 were selected as representative candidates for IgG-format validation. Sequencing showed that the three clones carry distinct CDR compositions, most divergent in HCDR3 and LCDR3, confirming non-redundant lineages rather than repeated isolates of a single clone (**Fig. 3A**).

To test whether the antigen reactivity observed in yeast display was retained in soluble format, all three clones were reformatted as chimeric IgG1 antibodies containing human IgG1 heavy-chain and human κ light-chain constant regions and titrated against immobilized Aβ_1-42_ AggreSure preparation by ELISA. All three bound in a concentration-dependent manner (**Fig. 3B**). CLAB17 showed the strongest apparent binding (apparent EC_50_ 23.5 ± 2.0 nM), followed by CLAB45 (84.7 ± 14.1 nM) and CLAB19 (116 ± 9 nM), corresponding to approximately 3.6-and 4.9-fold weaker apparent binding than CLAB17, respectively. Yeast-display reactivity was thus retained after IgG conversion, and the ELISA ranking defined CLAB17 and CLAB45 as parental candidates for humanization and CLAB19 as a weaker comparator that was not advanced.

An alternative Aβ-presenting probe, wtFc-Aβ_1-42_, was also tested in an antibody-coated ELISA; binding was weak and incompletely saturating over the tested range, precluding reliable apparent EC_50_ estimation, and this assay was not compared quantitatively with the immobilized-antigen ELISA (**Supplementary Fig. S2B,C**); the humanized variants behaved similarly (**Supplementary Fig. S3F**).

### Structure-guided design of focused humanization libraries for CLAB17 and CLAB45

CLAB17 and CLAB45 were selected for humanization because they combined confirmed IgG-format binding with distinct CDR architectures, allowing the focused humanization workflow to be evaluated in two antibody lineages. The goal was to reduce murine framework content while preserving the parental CDRs and detectable Aβ_1-42_AggreSure binding. Rather than committing to a single CDR-grafted design, we used a structure-guided, library-based strategy comprising CDR grafting onto human framework templates, structural comparison of parental and CDR-grafted Fv models, nomination of framework positions for focused back-mutation, humanization-library construction, single-round FACS screening, candidate selection, and IgG-format binding analysis (**Fig. 4A**). This design anticipates the well-documented risk that framework grafting perturbs CDR-loop conformation through vernier-zone and VH/VL packing residues, and treats that risk experimentally instead of assuming it away.

Parental murine and CDR-grafted humanized Fv models were generated and structurally superposed (**Fig. 4B,C**). The CDR-grafted CLAB17 model showed a larger whole-Fv deviation from its parental model than did CLAB45, with root-mean-square deviation (RMSD) values of 1.154 Å and 0.257 Å, respectively. These values were used only as qualitative heuristics for framework-position sampling.

Based on the structural models, a focused subset of framework positions was selected for back-mutation analysis. These positions were chosen from regions predicted to influence CDR support or VH/VL domain packing, but were not intended to represent an exhaustive set of all vernier-zone or packing residues. Focused humanization libraries were then built to sample the CDR-grafted human framework residue and the corresponding parental murine residue at these selected positions while preserving all parental CDR sequences (**Fig. 4D**; **Table 2**). This strategy converted humanization from a single fixed CDR-grafted design into a focused experimental search for framework combinations compatible with retained Aβ_1-42_ AggreSure binding.

### Single-round humanization screening recovered CLAB17-h2 and CLAB45-h8

Each focused humanization library was displayed on yeast and screened by FACS against the Aβ_1-42_ AggreSure probe, with Fab expression and antigen binding monitored simultaneously (**Fig. 5A**). This two-parameter readout allowed variants with poor Fab display to be distinguished from variants that displayed Fab but showed reduced antigen binding after framework grafting. Antigen-binding Fab-displaying populations were collected after one sorting round and used for single-clone isolation and characterization, rather than undergoing iterative rounds of library enrichment.

**Fig. 5.**
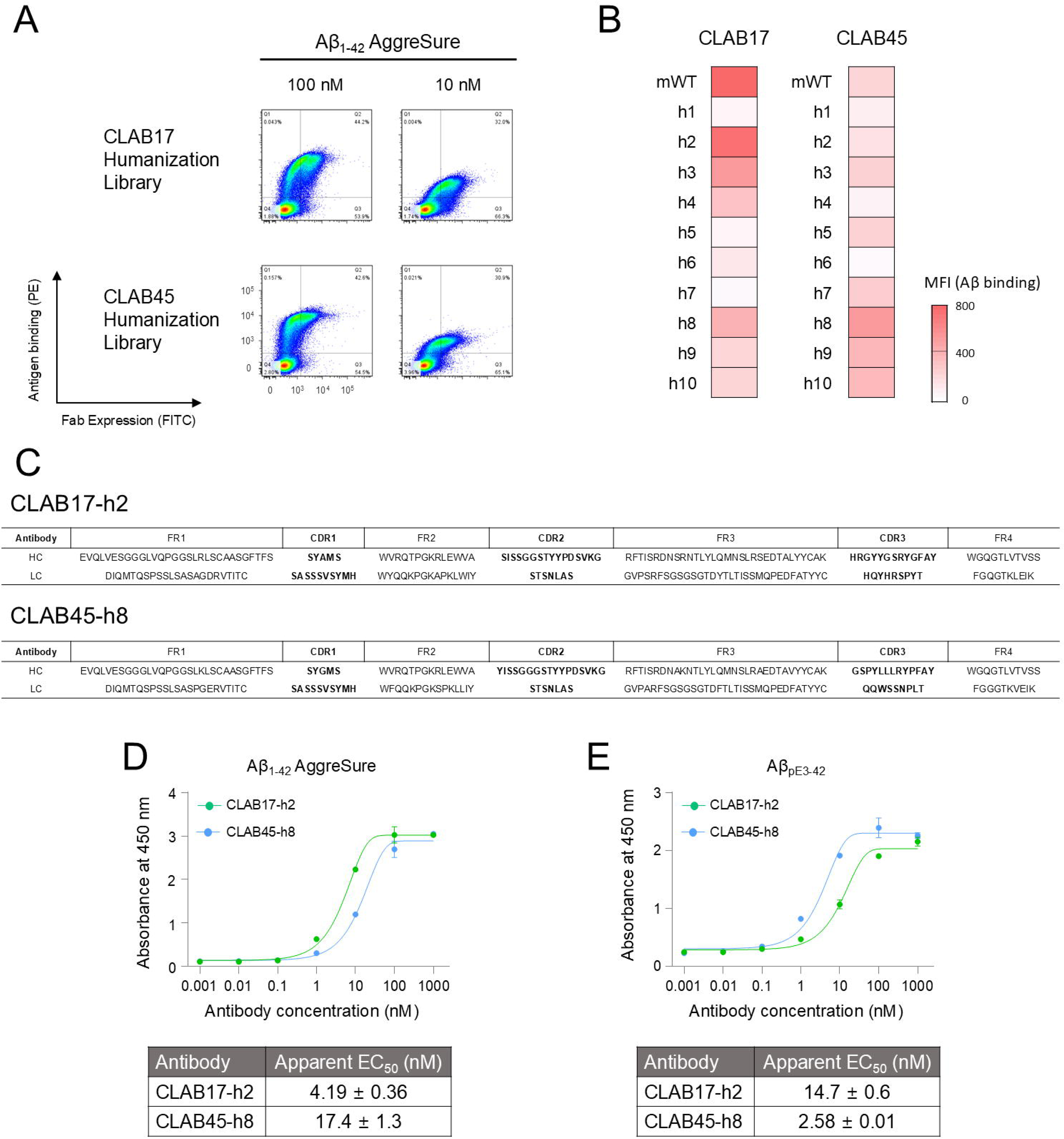
Single-round humanization screening identifies CLAB17-h2 and CLAB45-h8 as binding-positive variants. **(A)** Fluorescence-activated cell sorting (FACS) screening of CLAB17 and CLAB45 focused humanization libraries using amyloid-β_1-42_ (Aβ_1-42_) AggreSure preparation at 100 nM and 10 nM. Yeast-displayed fragment antigen-binding (Fab) variants were analyzed for Fab expression and antigen binding simultaneously, enabling collection of antigen-binding Fab-displaying populations after one sorting round. **(B)** Single-clone analysis of humanized variants h1–h10 and the murine parental reference (mWT) using 30 nM Aβ_1-42_ AggreSure preparation; antigen-binding mean fluorescence intensity (MFI) is shown. CLAB17-h2 and CLAB45-h8 were selected as representative binding-positive variants for immunoglobulin G (IgG)-format validation based on single-clone FACS profiles. **(C)** Final sequences of CLAB17-h2 and CLAB45-h8 by framework and complementarity-determining region (CDR); both retained all parental CDRs within humanized frameworks. **(D)** Binding of IgG-formatted CLAB17-h2 and CLAB45-h8 to immobilized Aβ_1-42_ AggreSure preparation. Apparent half-maximal effective concentration (EC_50_) values were 4.19 ± 0.36 nM for CLAB17-h2 and 17.4 ± 1.3 nM for CLAB45-h8. **(E)** Binding of IgG-formatted CLAB17-h2 and CLAB45-h8 to immobilized pyroglutamate-modified amyloid-β_3-42_ (Aβ_pE3-42_). Apparent EC_50_ values were 14.7 ± 0.6 nM for CLAB17-h2 and 2.58 ± 0.01 nM for CLAB45-h8. Apparent EC_50_ values are reported as descriptive enzyme-linked immunosorbent assay (ELISA) readouts under each immobilized-antigen assay condition and should not be interpreted as monovalent affinity constants or as definitive evidence of intrinsic antigen preference. Data are shown as mean ± standard deviation (SD).

Ten clones per lineage were then isolated and characterized by single-clone FACS (**Fig. 5B** and **Supplementary Fig. S3A**). Candidate variants were triaged using single-clone FACS profiles, including antigen-binding signal and qualitative Fab-display behavior. CLAB17-h2 and CLAB45-h8 were selected as representative binding-positive variants for IgG-format validation. Recovery of binding-positive variants from both lineages after one FACS sorting round indicates that the focused humanization libraries sampled framework combinations compatible with detectable Aβ_1-42_ AggreSure binding under the tested screening condition.

Both final variants preserved the complete parental CDR set within humanized frameworks (**Fig. 5C**), with no CDR-region changes introduced during humanization. Position-resolved comparison with the CDR-grafted human and parental murine frameworks showed lineage-dependent retention: among 24 sampled positions CLAB17-h2 retained 14 human and 9 parental murine residues (6 VH, 3 VL) plus one off-design residue at VL Kabat 15 (alanine, distinct from both the human valine and the murine proline), whereas among 20 sampled positions CLAB45-h8 retained 9 human and 11 parental murine residues (5 VH, 6 VL). This lineage-dependent pattern supports experimental sampling of framework combinations rather than assuming uniform tolerance of a fixed CDR-grafted framework. **Table 2** reports residue identity directly from the final sequenced variants rather than inferring human-framework retention from unannotated positions.

### Humanization increased predicted humanness while retaining binding; final variants showed favorable in silico developability triage profiles

Because the value of humanization lies in reducing murine sequence content while preserving binding, we quantified predicted humanness by comparing the parental murine antibodies with the final selected humanized variants. CDR-grafted sequences were treated as design intermediates for focused library construction and were not included in the final humanness comparison. Across both lineages, the final humanized variants showed increased overall OASis scores compared with their parental murine antibodies. For CLAB17, the overall OASis score increased from 0.659 in the parental murine antibody to 0.825 in CLAB17-h2. For CLAB45, the overall OASis score increased from 0.646 in the parental murine antibody to 0.811 in CLAB45-h8.

This increase was driven primarily by improved light-chain humanness. VL OASis identity increased from 0.398 to 0.765 for CLAB17 and from 0.408 to 0.755 for CLAB45, whereas VH OASis identity remained high in both lineages. Independent humanness metrics showed the same overall trend. Hu-mAb VH/VL scores increased from 0.000/0.000 to 0.385/0.765 (CLAB17-h2) and 0.430/0.615 (CLAB45-h8), and T20 scores from 77.9/63.7 to 85.5/82.1 and from 77.8/65.2 to 83.4/80.1, respectively. The final humanized variants were assigned to human germline genes IGHV3-23*04 and IGKV1-39*01, with germline identities of 90.8% and 82.1% for CLAB17-h2 VH and VL, respectively, and 89.8% and 80.0% for CLAB45-h8 VH and VL, respectively (**Table 1**). Together, these results show increased predicted humanness across two independent lineages while preserving parental CDRs and Aβ-preparation binding; the scores are interpreted as relative improvements over the parental sequences.

To triage early developability without committing extensive bench resources, we scanned the final humanized sequences in silico for common sequence liabilities and applied TAP to the modeled Fv structures. Neither CLAB17-h2 nor CLAB45-h8 contained extra cysteines beyond the conserved VH/VL intradomain disulfide-forming cysteines, and no N-linked glycosylation sequons, CDR-resident NG/NS deamidation-associated motifs, or CDR-resident DG isomerization-associated motifs were detected. CDR-resident methionine residues were present in HCDR1 and LCDR1 of both antibodies, and CLAB45-h8 additionally contained a tryptophan residue in LCDR3; these residues were recorded as potential oxidation-monitoring sites.

TAP placed both humanized variants within the guideline ranges for all five evaluated metrics, and neither received an amber or red flag (**Supplementary Table S2**). Experimentally, CLAB17-h2 and CLAB45-h8 were expressed in transient HEK293F culture at 80.3 and 48.3 mg/L, respectively. ImageJ densitometry of Coomassie-stained SDS-PAGE gels estimated purities of >99% for CLAB17-h2 and approximately 96.5% for CLAB45-h8. Both antibodies migrated predominantly as intact IgG species under non-reducing conditions and showed the expected heavy-and light-chain bands under reducing conditions (**Supplementary Fig. S3B**). Together, these analyses support early-stage computational and gross protein-quality triage, but not comprehensive experimental developability qualification.

### Humanized CLAB17-h2 and CLAB45-h8 show distinct Aβ-preparation binding profiles

Having confirmed detectable binding and increased predicted humanness, we compared the two humanized lineages across the tested Aβ preparations. In the immobilized Aβ_1-42_AggreSure ELISA, CLAB17-h2 and CLAB45-h8 showed apparent EC_50_ values of 4.19 ± 0.36 nM and 17.4 ± 1.3 nM, respectively (**Fig. 5D**). In the immobilized Aβ_pE3-42_ELISA, the corresponding values were 14.7 ± 0.6 nM and 2.58 ± 0.01 nM (**Fig. 5E**). Thus, CLAB17-h2 showed an approximately 3.5-fold lower apparent EC_50_ on Aβ_1-42_ AggreSure, whereas CLAB45-h8 showed an approximately 6.7-fold lower apparent EC_50_ on Aβ_pE3-42_. Because these immobilized preparations may differ in assembly state, coating efficiency, epitope density, and orientation, these differences are not interpreted as intrinsic antigen preference or pyroglutamate-specific recognition.

In exploratory non-target binding controls, biotinylated SARS-CoV-2 spike receptor-binding domain (RBD), programmed cell death protein 1 (PD-1), and albumin were tested at 100 nM with Aβ_1-42_ AggreSure as the target-positive control (**Supplementary Fig. S3C–E**). With the Aβ_1-42_ AggreSure signal normalized to 100%, CLAB17-h2 showed relative signals of 8.04 ± 0.32%, 3.29 ± 0.08%, and 3.53 ± 0.16% for RBD, PD-1, and albumin, respectively, whereas CLAB45-h8 showed 17.75 ± 0.98%, 9.28 ± 2.10%, and 9.15 ± 1.31%. Because the assay lacked background subtraction, irrelevant IgG controls, and matched biotin-labeling ratios, these measurements are reported as exploratory non-target binding controls rather than a quantitative polyspecificity assessment.

Overall, the workflow recovered humanized variants from two lineages that showed detectable apparent Aβ-preparation binding while showing increased predicted humanness and distinct ELISA reactivity profiles. Because parental and humanized antibodies were assayed on separate plates, the lower apparent EC_50_ values of the humanized variants are not interpreted as affinity improvements conferred by humanization.

## Discussion

This study establishes an integrated immune yeast Fab-display and single-round focused humanization pipeline and demonstrates it by generating binding-positive humanized antibodies across two sequence-distinct anti-Aβ lineages. The principal advance is methodological: an antigen-experienced murine repertoire was coupled directly to focused framework sampling, allowing humanized variants to be recovered after a single humanization sorting round while preserving parental CDRs and improving predicted humanness. Because the Aβ-derived preparations were not biophysically resolved into defined assembly states, the antibodies are described as Aβ-preparation-binding candidates; humanness and developability metrics are used as engineering triage rather than therapeutic qualification.

Anti-Aβ antibodies have been generated by hybridoma screening (Sehlin et al., 2010; Ying et al., 2009), phage and other display libraries (Frenzel et al., 2016; Manoutcharian et al., 2003; Medecigo et al., 2010), human memory B-cell cloning (Budd Haeberlein et al., 2022; Sevigny et al., 2016), human-transgenic mice (Green, 2014; Lonberg, 2008), and rational selection against defined Aβ species (Antonios et al., 2015; Mintun et al., 2021). Immune yeast Fab-display combines in vivo affinity maturation with simultaneous measurement of Fab display and antigen binding, which is useful for heterogeneous or self-associating antigens whose valency and presentation can distort binding signals. Subsequent IgG-format validation provided an orthogonal triage step; CLAB19, for example, was recovered during display screening but not advanced because of weaker validated binding.

Using a murine immune repertoire provides a robust source of affinity-matured binders but requires subsequent framework engineering. We addressed this with focused humanization libraries rather than a single fixed CDR-grafted design. Yeast display has been applied to antibody humanization before, for example to humanize chicken-derived single-chain variable fragments by combining framework libraries with sequencing-guided selection over successive sorting rounds (Elter et al., 2021). The present workflow differs in starting from a murine immune Fab repertoire, restricting the design to a structure-nominated binary framework set, and recovering variants from two independent lineages after a single sorting round. Structural models nominated framework positions predicted to influence CDR support or VH/VL packing, and library-based screening recovered binding-positive variants from both lineages. OASis, Hu-mAb, and T20 scores increased after humanization, while final sequence analysis documented lineage-dependent retention of parental framework residues (**Tables 1 and 2**). CLAB17-h2 retained human identities at most sampled positions, whereas CLAB45-h8 retained parental murine residues at slightly more than half, underscoring that framework tolerance is lineage dependent. Thus, the practical strength of the workflow lies in combining structure-guided nomination with empirical selection of binding-compatible framework combinations.

The humanized antibodies also showed different apparent ELISA reactivity profiles: CLAB17-h2 had a lower apparent EC_50_ on Aβ_1-42_ AggreSure, whereas CLAB45-h8 had a lower apparent EC_50_ on Aβ_pE3-42_. These differences support retaining both variants for further characterization but do not establish intrinsic antigen preference, pyroglutamate-specific recognition, or assembly-state selectivity because the assays used bivalent IgG and immobilized, heterogeneous antigen preparations. Defining the molecular basis of these profiles will require fractionated or structurally defined Aβ species, matched unmodified and pyroglutamate-modified peptides, coating-independent competition assays, and monovalent Fab-based SPR or BLI (Söderberg et al., 2023; Talucci et al., 2026).

Other secondary assays are similarly interpreted within their intended scope. The wtFc-Aβ_1-42_ binding profiles were not quantifiable, the non-target panel was exploratory rather than a formal polyspecificity assay, and the sequence-liability/TAP analyses represent computational developability triage rather than experimental qualification. These limitations do not affect the central workflow result that focused humanization recovered binding-positive variants from two independent lineages, but they define the next stage of candidate characterization. Future studies should determine which molecular Aβ species are recognized and whether the observed ELISA profiles reflect intrinsic affinity, avidity, epitope density, or immobilization effects. Biophysical characterization of the antigen preparations, binding assays using defined Aβ species, monovalent SPR/BLI, staining of AD and non-AD brain tissue, comparison with established Aβ antibodies, and cellular functional assays would address these questions (Jain et al., 2017; Söderberg et al., 2023; Talucci et al., 2026). At present, CLAB17-h2 and CLAB45-h8 are best positioned as humanized Aβ-preparation-binding reagents with increased predicted humanness and favorable computational triage profiles. More broadly, the workflow provides a reusable route from immune yeast-display discovery to single-round focused humanization for complex, self-associating antigens.

## Supporting information

Graphical abstract caption

## Acknowledgments

This work was supported in part by the Innovative New Drug Development Program funded by the Ministry of Science and ICT (MSIT) (grant number RS-2026-25527526 to C.-H.L.), the Global Physician-Scientist Program funded by the Ministry of Health and Welfare (MOHW) (grant number RS-2025-25459535 to C.-H.L.), and the Mid-Career Researcher Program funded by MSIT (grant number RS-2023-00278980 to C.-H.L.).

## Ethics Approval

All animal experiments were conducted in accordance with institutional guidelines for the care and use of laboratory animals. The animal study protocol was approved by the Institutional Animal Care and Use Committee of Seoul National University under protocol number SNU-230519-9-3. No human participants or human-derived specimens were used in this study.

## Author Contributions

Y.K. designed and performed experiments, analyzed data, and wrote the manuscript. H.K., J.S., and Y.L. contributed to the construction of antibody and antigen expression vectors and protein purification. M.P. contributed to in silico humanness, IgBLAST-based germline identity, and developability analyses of the humanized antibodies. C.-H.L. conceived and supervised the study, secured funding, interpreted data, and revised the manuscript.

## Data Availability Statement

All data that support the findings of this study are available from the corresponding author upon reasonable request.

## Supplementary information

**Supplementary Fig. S1. Single-clone FACS of immune-library-derived Fab clones.**

**Supplementary Fig. S2. SDS-PAGE and wtFc-Aβ_1-42_ binding analysis of parental CLAB antibodies and the wtFc-Aβ_1-42_ fusion probe.**

**Supplementary Fig. S3. Single-clone analysis, IgG quality control, non-target protein quality control, non-target protein binding, and wtFc-Aβ_1-42_ binding of humanized CLAB antibodies.**

**Supplementary Table S1. Primers for murine VH and VL amplification.**

**Supplementary Table S2. In silico developability liability scan of humanized CLAB17-h2 and CLAB45-h8.**

**Full legends, Supplementary Methods, and Supplementary Tables S1 and S2 are provided in the Supplementary information file.**

---

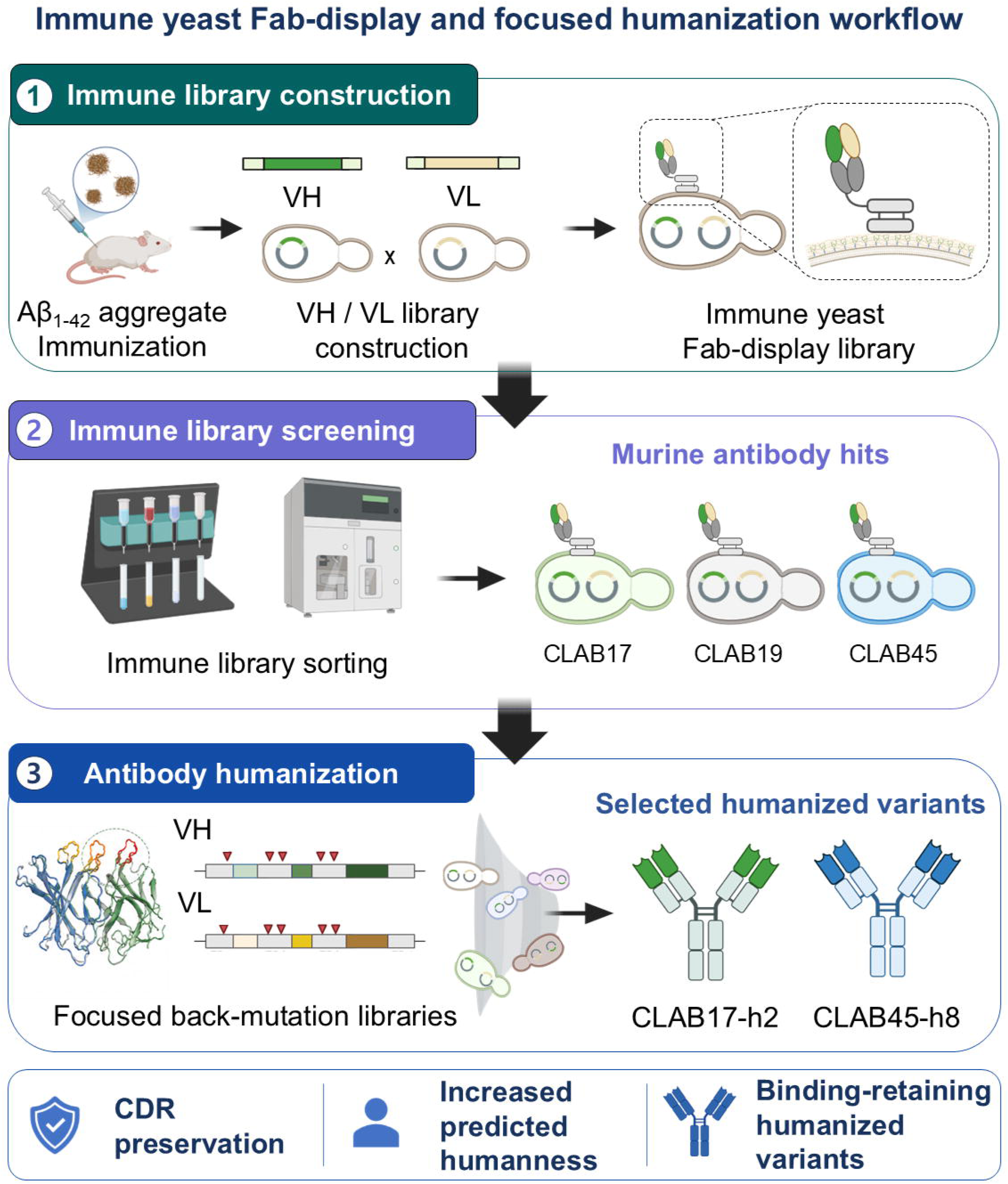

