## Supplementary material for "Focused framework sampling recovers binding-positive humanized anti-amyloid-β antibodies in a single sorting round": Graphical abstract caption

An immune yeast Fab-display library from Aβ₁₋₄₂-immunized mice was screened by MACS and FACS to recover three antibodies. Structure-guided libraries sampling selected framework positions yielded humanized CLAB17-h2 and CLAB45-h8 after a single sorting round, preserving parental CDRs and showing increased predicted humanness.
