## Supplementary information for "Focused framework sampling recovers binding-positive humanized anti-amyloid-β antibodies in a single sorting round"

**Supplementary Methods**

**Preparation of Aβ_1-42_ aggregate preparations**

Amyloid-β_1-42_ (Aβ_1-42_) aggregate preparations were generated as previously described (Stine et al., 2011), with minor modifications. Lyophilized Aβ_1-42_ peptide (AnaSpec, Fremont, CA, USA; AS-20276) was dissolved in hexafluoroisopropanol (HFIP) to 1 mM, incubated at room temperature for 30 min, and evaporated overnight in a chemical fume hood. Residual HFIP and moisture were removed by drying the resulting film for 1 h in a HyperVAC-LITE vacuum concentrator (Hanil Scientific Inc., Gimpo, Republic of Korea; HVC-2124). The dried film was dissolved in dimethyl sulfoxide (DMSO) at 5 mg/mL, diluted into phosphate-buffered saline (PBS, pH 7.4) to 100 μM, and incubated at 37 °C for 7 days. This material was used for mouse immunization and serum enzyme-linked immunosorbent assay (ELISA). Preparations were not further fractionated into defined monomeric, oligomeric, protofibrillar, or fibrillar species; binding data were therefore interpreted as reactivity to the tested Aβ_1-42_-derived preparations.

**Mouse immunization and serum collection**

Mouse immunization was performed as previously described, with minor modifications (Chung et al., 2026; Lim et al., 2026). Five six-week-old BALB/c mice were immunized intraperitoneally with 50 μg of Aβ_1-42_ aggregate preparation mixed at a 1:1 volume ratio with alhydrogel adjuvant (InvivoGen, San Diego, CA, USA; vac-alu-50). Five immunizations were administered at 2-week intervals (weeks 1, 3, 5, 7, and 9), and blood was collected during the course to monitor the serum antibody response. Serum was separated by centrifugation and stored at -80 °C until analysis. All animal procedures followed institutional guidelines and were approved by the Institutional Animal Care and Use Committee of Seoul National University (SNU-230519-9-3).

**General ELISA procedure**

All ELISAs were performed on 96-well MaxiSorp microplates (Thermo Fisher Scientific, Waltham, MA, USA; 439454) as previously described, with minor modifications (Kang et al., 2024; Yang et al., 2026). Plates were coated overnight at 4 °C, blocked with 3% bovine serum albumin (BSA; Sigma-Aldrich, St. Louis, MO, USA) in PBS for 1 h at room temperature, and washed three times with PBS containing 0.05% Tween-20 (PBST). Samples and detection reagents were each incubated for 1 h at room temperature, with three PBST washes (300 μL per well) between steps. Color was developed with 3,3′,5,5′-tetramethylbenzidine (TMB; Thermo Fisher Scientific), the reaction was stopped with 2 M H2SO4, and absorbance at 450 nm was measured on an Infinite 200 PRO NanoQuant microplate reader (Tecan Trading AG, Männedorf, Switzerland). Assay-specific coating material, sample dilutions, and detection reagents are given below.

**Serum ELISA**

Aβ_1-42_ aggregate preparation was coated at 100 ng per well. Mouse serum was serially diluted 10-fold in blocking buffer, and bound antibody was detected with horseradish peroxidase (HRP)-conjugated goat anti-mouse immunoglobulin G (IgG) (heavy and light chains, H+L) secondary antibody (1:8,000; Invitrogen, Carlsbad, CA, USA; 31430).

**Construction of the immune yeast Fab-display library**

Following the final boost, mice were euthanized and spleen and bone marrow were collected for immune repertoire recovery. Messenger RNA was isolated and complementary DNA (cDNA) synthesized with the First Strand cDNA Synthesis Kit (Thermo Fisher Scientific; 18080400). Heavy-chain variable (VH) and light-chain variable (VL) genes were amplified with murine antibody-specific primer sets (**Supplementary Table S1**) and fused to the first heavy-chain constant domain (CH1) and the light-chain constant region (CL), respectively, by overlap polymerase chain reaction (PCR).

Library transformation, culture, and mating followed the yeast fragment antigen-binding (Fab)-display platform described previously (Kim et al., 2026). Briefly, 4 μg of VH-CH1 fragment was co-transformed with 12 μg of linearized pYDS-H into Saccharomyces cerevisiae JAR200 (MATa), and 4 μg of VL-CL fragment with 12 μg of linearized pYDS-K into S. cerevisiae YvH10 (MATα), by electroporation-mediated homologous recombination. Because pYDS-H carries TRP1 and pYDS-K carries URA3, the JAR200 heavy-chain library was maintained in 300 mL of synthetic dextrose casamino acid (SDCAA) medium supplemented with uracil and the YvH10 light-chain library in SDCAA supplemented with tryptophan (30 °C, 160 rpm, 15 h, to an optical density at 600 nm [OD_600_] of approximately 5). Colony diversity of each haploid library was determined by serial dilution and plating on the corresponding selective agar (**Fig. 1D**).

For mating, heavy-chain and light-chain library cells were adjusted to OD_600_ = 100, mixed at equal volumes, plated on yeast extract-peptone-dextrose (YPD) agar at pH 4.5, and incubated at 30 °C for 8 h. Mated cells were collected, washed twice with sterile distilled water, resuspended in SDCAA at OD_600_ = 0.1, and cultured overnight at 30 °C and 160 rpm. Because growth on unsupplemented SDCAA requires complementation of both TRP1 and URA3, colonies on this medium report diploids, whereas tryptophan-supplemented SDCAA additionally permits growth of URA3-positive YvH10-derived cells; mating efficiency was calculated as the ratio of the two colony counts (**Fig. 1D**).

**Biotinylated Aβ-derived probes for yeast display screening**

Two biotinylated amyloid-β (Aβ)-derived probes were prepared in house so that enrichment could be monitored consistently across selection rounds. The Aβ_1-42_ aggregate probe was generated by subjecting N-terminally biotinylated Aβ_1-42_ peptide (AnaSpec; AS-23524-01) to the aggregation procedure described above. The Aβ_1-42_ AggreSure probe was generated by labeling Aβ_1-42_ AggreSure (AnaSpec; AS-72216) with EZ-Link Sulfo-NHS-Biotin (Thermo Fisher Scientific; 21217) per the manufacturer's instructions, followed by removal of free biotin with Zeba Dye and Biotin Removal Spin Columns (Thermo Fisher Scientific; A44298).

**Screening and sorting of the yeast-displayed Fab library**

Induction, surface staining, and sorting were performed as previously described (Kim et al., 2026). Induced cells were incubated with mouse anti-FLAG antibody (1:200; Thermo Fisher Scientific; MA1-91878), which detects the C-terminal FLAG epitope fused to the light chain, and the indicated biotinylated probe for 1 h at room temperature, washed with PBS containing 0.1% BSA (PBSA, pH 7.4), and stained with fluorescein isothiocyanate (FITC)-conjugated anti-mouse IgG (1:200; Thermo Fisher Scientific; 62-6511) and streptavidin-phycoerythrin (PE) (1:200; Invitrogen; S866) for 15 min at 4 °C in the dark. Singlets were defined by diagonal gating on forward scatter area versus height. For analytical profiles, each probe was used at 100 and 10 nM (**Fig. 2A**).

For magnetic-activated cell sorting (MACS), approximately 1 × 10^9^ induced cells were incubated with 100 nM biotin-Aβ_1-42_ aggregate probe for 1 h at room temperature, washed with PBSA, resuspended with streptavidin-conjugated magnetic microbeads (Miltenyi Biotec, Bergisch Gladbach, Germany; 130-048-102), and applied to an LS column (Miltenyi Biotec; 130-042-401) on a MidiMACS separator. The column was washed five times with 10 mL of PBSA, and retained cells were eluted with 6 mL of SDCAA and expanded overnight at 30 °C. Fluorescence-activated cell sorting (FACS) was then performed on an SH800S cell sorter (Sony Biotechnology Inc., San Jose, CA, USA) using 50 nM biotin-Aβ_1-42_ aggregate probe, collecting the top 1% antigen-binding population within the FITC-positive Fab-expressing gate. The pre-sort, MACS-enriched, and post-FACS populations were designated R1, R2, and R3, respectively.

R3 cells were expanded in SDCAA (30 °C, 160 rpm, 24 h) and plated on SDCAA agar. Individual colonies were cultured in 3 mL of SDCAA (30 °C, 160 rpm, 24 h) and induced in 2× synthetic galactose casamino acid (SGCAA) medium at an initial OD_600_ of 0.5 (20 °C, 160 rpm, 48 h). Induced clones were stained separately with 30 nM of each biotinylated probe, and clones binding both were prioritized (**Fig. 2B**). VH and VL sequences were determined by colony PCR and sequence analysis. Enrichment was reported as the fraction in the upper-right quadrant (Q2) of the displayed analysis, corresponding to FITC-positive/PE-positive events among gated singlets at R1-R3 (**Fig. 2A**); gates were set with unstained and single-stain controls, and this definition was applied consistently to the values reported in the Results and figure legend.

**Expression and purification of selected antibodies**

Antibodies were expressed and purified as previously described (Lim et al., 2026). VH and VL fragments were subcloned by Gibson assembly into pCIW-VH and pCIW-VL vectors encoding the human IgG1 heavy-chain and human kappa light-chain constant regions, and the plasmids were co-transfected into human embryonic kidney 293F (HEK293F) cells at a heavy-chain:light-chain ratio of 1:3. Five days later, supernatants were harvested and antibodies captured on Protein A resin (Amicogen, Jinju, Republic of Korea; Puriose ProA FF-30), washed with 50 mL of 1× PBS, eluted with 12 mL of 100 mM glycine (pH 2.5), and immediately neutralized with 1 mL of 1 M Tris-HCl (pH 8.0). Buffer exchange into PBS (pH 7.4) used Amicon Ultra-4 30-kDa centrifugal filters (Merck Millipore). Purified antibodies were analyzed by 12% sodium dodecyl sulfate-polyacrylamide gel electrophoresis (SDS-PAGE) (**Supplementary Fig. S2A and S3B**), with purity estimated by ImageJ densitometry of Coomassie-stained bands.

**ELISA against immobilized Aβ-derived preparations**

Aβ_1-42_ AggreSure or pyroglutamate-modified Aβ_3-42_ (Aβ_pE3-42_; AnaSpec; AS-29907-01) was coated at 2 μg/mL (50 μL per well). Serially diluted antibodies were applied, and bound IgG was detected with HRP-conjugated anti-human IgG fragment crystallizable (Fc) antibody (1:12,000; Arigo Biolaboratories, Hsinchu, Taiwan; ARG23874).

**Preparation of recombinant wtFc-Aβ_1-42_ and non-target proteins**

The wild-type Fc (wtFc)-Aβ_1-42_ coding sequence was cloned into pcDNA3.4 by Gibson assembly to generate an Fc-mediated Aβ_1-42_-presenting fusion. Non-target proteins, comprising the SARS-CoV-2 spike receptor-binding domain (RBD; Wuhan-Hu-1), human programmed cell death protein 1 (PD-1), and human albumin, were cloned into the same vector with a polyhistidine (His) tag. All proteins were expressed in Expi293 cells (Thermo Fisher Scientific); wtFc-Aβ_1-42_ was purified on Protein A resin and His-tagged proteins on nickel-nitrilotriacetic acid (Ni-NTA) agarose, followed by buffer exchange into PBS. Preparations were analyzed by SDS-PAGE under reducing and non-reducing conditions (**Supplementary Fig. S2B and S3C**). Purified proteins were then biotinylated with EZ-Link Sulfo-NHS-Biotin and freed of unreacted biotin as described above.

**Antibody-coated ELISA for wtFc-Aβ_1-42_ and non-target proteins**

Antibodies were coated at 2 μg/mL (50 μL per well). For wtFc-Aβ_1-42_ binding, serially diluted biotinylated wtFc-Aβ_1-42_ was applied. For the non-target panel, biotinylated RBD, PD-1, and albumin were tested at 100 nM alongside biotinylated Aβ_1-42_ AggreSure as the target-positive control. Bound probes were detected with HRP-conjugated streptavidin (1:40,000; Abcam, Cambridge, MA, USA; ab7403). For normalized non-target analysis, the raw optical density at 450 nm (OD_450_) for each non-target protein was divided by the raw OD_450_ obtained with biotinylated Aβ_1-42_ AggreSure for the same antibody, without background subtraction, with the Aβ_1-42_ AggreSure signal set to 100%.

**Design and screening of focused humanization libraries**

For each selected murine antibody, the parental heavy-chain and light-chain variable regions were compared with human germline V genes, and the germline with the highest sequence identity was used as the human acceptor framework. The parental complementarity-determining regions (CDRs) were then transferred onto this framework by CDR grafting. Variable fragment (Fv) models of the parental murine and CDR-grafted antibodies were generated using AlphaFold 3 and structurally compared (**Fig. 4B,C**). Based on these models, a focused subset of framework positions was selected for back-mutation analysis from regions predicted to influence CDR support or VH/VL domain packing; this subset was not intended to be an exhaustive set of vernier-zone or packing residues. Each selected position was sampled between the CDR-grafted human residue and the corresponding parental murine residue, with all parental CDR sequences preserved (**Fig. 4D; Table 2**).

Designed VH and VL fragments were introduced into yeast and mated to generate paired Fab humanization libraries using the strategy described above for the immune library (Kim et al., 2026). Libraries were induced in 2× SGCAA at an initial OD_600_ of 0.5 (20 °C, 160 rpm, 48 h) and stained as described above with biotinylated Aβ_1-42_ AggreSure at 100 or 10 nM. Fab expression and antigen binding were monitored simultaneously on an SH800S cell sorter, and antigen-binding Fab-displaying populations were collected after a single sorting round (**Fig. 5A**).

Sorted cells were streaked directly onto SDCAA agar for single-clone isolation. For each lineage, 10 humanized candidates were analyzed for Fab expression and binding to 30 nM Aβ_1-42_ AggreSure by single-clone FACS. Candidates were triaged using antigen-binding signal and qualitative Fab-display behavior, and representative binding-positive variants were selected for IgG-format validation, yielding CLAB17-h2 and CLAB45-h8 (**Fig. 5B,C**). Final variants were sequenced to record any off-design residues arising during library construction, amplification, or selection.

**Humanness and germline analysis**

Humanness of the parental murine antibodies and the final humanized variants was evaluated using the BioPhi OASis (Observed Antibody Space identity search), Hu-mAb, and T20 humanness scores. Germline assignments were determined by Immunoglobulin BLAST (IgBLAST) against species-appropriate international ImMunoGeneTics information system (IMGT) germline databases: parental murine antibodies against mouse germline genes and final humanized variants against human germline genes. CDR-grafted sequences served as design intermediates and were excluded from the final comparison because they were not selected antibody products (**Table 1**).

**Sequence-based developability assessment**

Final humanized VH/VL sequences were scanned in silico for developability liabilities, including extra or unpaired cysteines, N-linked glycosylation sequons (N-X-S/T), deamidation (NG, NS) and isomerization (DG) motifs, and methionine/tryptophan oxidation, with particular attention to CDR-resident motifs. The Therapeutic Antibody Profiler (TAP) was applied to the Fv models to evaluate total CDR length, CDR-vicinity patches of surface hydrophobicity (PSH), positive charge (PPC), and negative charge (PNC), and the structural Fv charge symmetry parameter (SFvCSP) (Raybould et al., 2019). Amber and red flags were interpreted according to the clinical-stage therapeutic antibody guideline thresholds (**Supplementary Table S2**).

**Scope and interpretation of binding and developability assays**

Apparent half-maximal effective concentration (EC_50_) values were derived from ELISA dose-response curves and interpreted as assay-dependent, avidity-influenced readouts rather than monovalent affinity constants. Because the wtFc-Aβ_1-42_ and non-target assays used an antibody-coated, biotinylated-probe format, the resulting data were interpreted separately from the immobilized-antigen ELISAs and were not compared quantitatively across assay formats. The degree of biotin labeling was not independently determined, so signals were not compared across probes with matched labeling stoichiometry. SDS-PAGE served as a gross quality check and was not intended to establish oligomeric stoichiometry, biotin-labeling ratio, conformational integrity, or functional equivalence. Sequence liabilities and TAP flags are computational triage results, not experimentally confirmed liabilities.

**Data analysis and software**

Where applicable, quantitative data are presented as mean ± standard deviation (SD). No formal statistical significance testing was performed. ELISA dose-response curves and apparent EC_50_ values were calculated using the EC_50_ calculator tool provided by AAT Bioquest (AAT Bioquest, Inc., Sunnyvale, CA, USA). Graphs were generated using GraphPad Prism 9 (GraphPad Software, San Diego, CA, USA). Schematic illustrations, including the graphical abstract, **Fig. 1A**, and **Fig. 4A**, were created using BioRender.com.

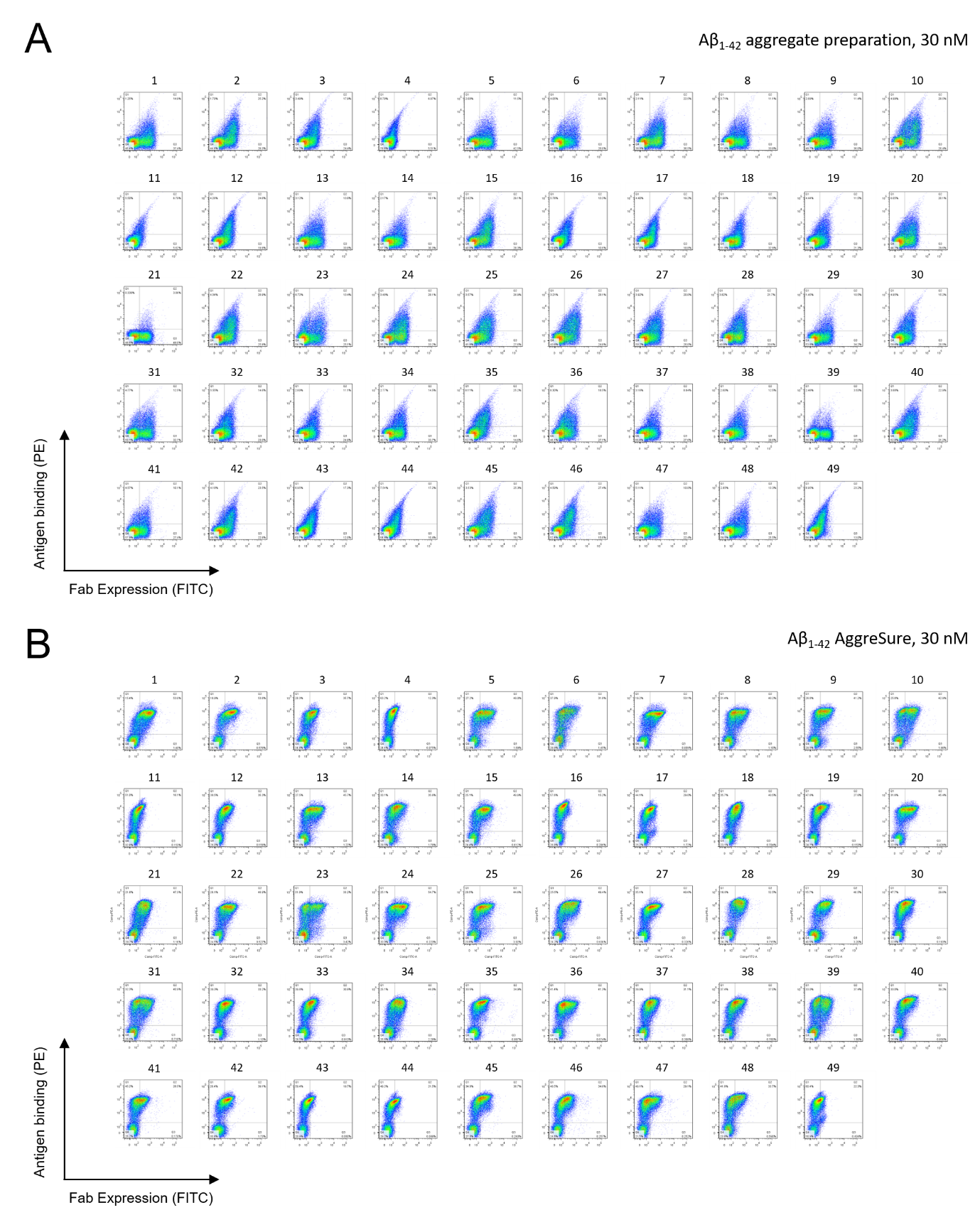

**Supplementary Fig. S1. Single-clone fluorescence-activated cell sorting (FACS) of immune-library-derived fragment antigen-binding (Fab) clones.** FACS profiles of all 49 individual yeast-displayed Fab clones analyzed after immune-library screening. **(A)** Amyloid-β_1-42_ (Aβ_1-42_) aggregate probe at 30 nM. **(B)** Aβ_1-42_ AggreSure probe at 30 nM. Fab expression, fluorescein isothiocyanate (FITC); antigen binding, phycoerythrin (PE). Used to support clone triage together with immunoglobulin G (IgG)-format enzyme-linked immunosorbent assay (ELISA) validation.

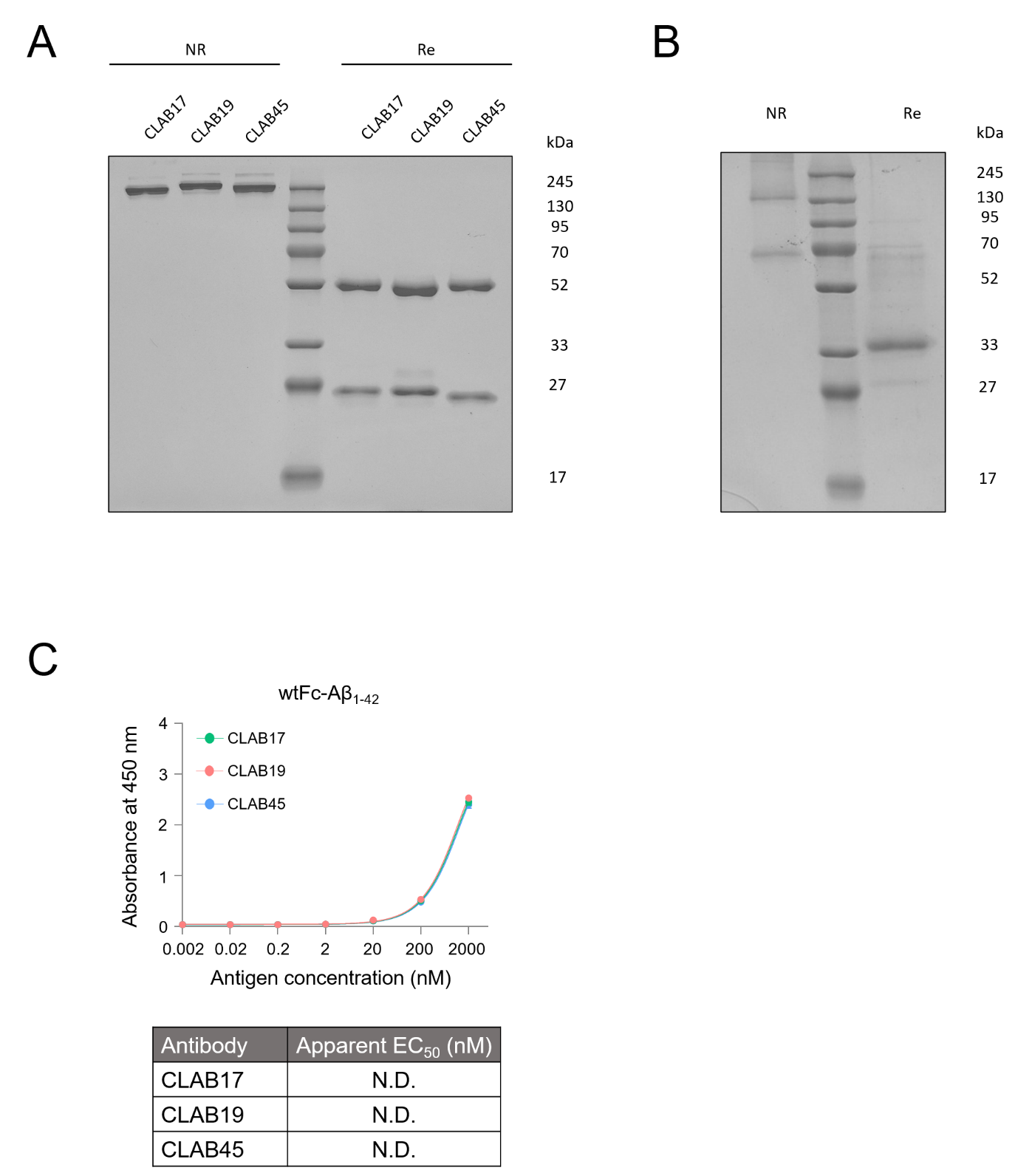

**Supplementary Fig. S2. Sodium dodecyl sulfate-polyacrylamide gel electrophoresis (SDS-PAGE) and wild-type Fc (wtFc)-Aβ_1-42_ binding analysis of parental CLAB antibodies.** **(A)** Reducing (Re) and non-reducing (NR) SDS-PAGE of purified immunoglobulin G (IgG)-formatted CLAB17, CLAB19, and CLAB45; markers in kDa. **(B)** SDS-PAGE of purified wtFc-Aβ_1-42_. Two major non-reducing species at approximately 70 and 130 kDa collapsed under reducing conditions into a single band near 30–33 kDa, indicating reducing-sensitive higher-order species; SDS-PAGE alone does not establish oligomeric stoichiometry. **(C)** Antibody-coated enzyme-linked immunosorbent assay (ELISA) of CLAB17, CLAB19, and CLAB45 against serially diluted biotinylated wtFc-Aβ_1-42_ (0.002–2000 nM), a fragment crystallizable (Fc)-mediated Aβ_1-42_-presenting probe. Binding was weak and incompletely saturating over the tested range, so apparent half-maximal effective concentration (EC_50_) values were not determined (N.D.); this does not indicate absence of wtFc-Aβ_1-42_ binding. Data are mean ± standard deviation (SD).

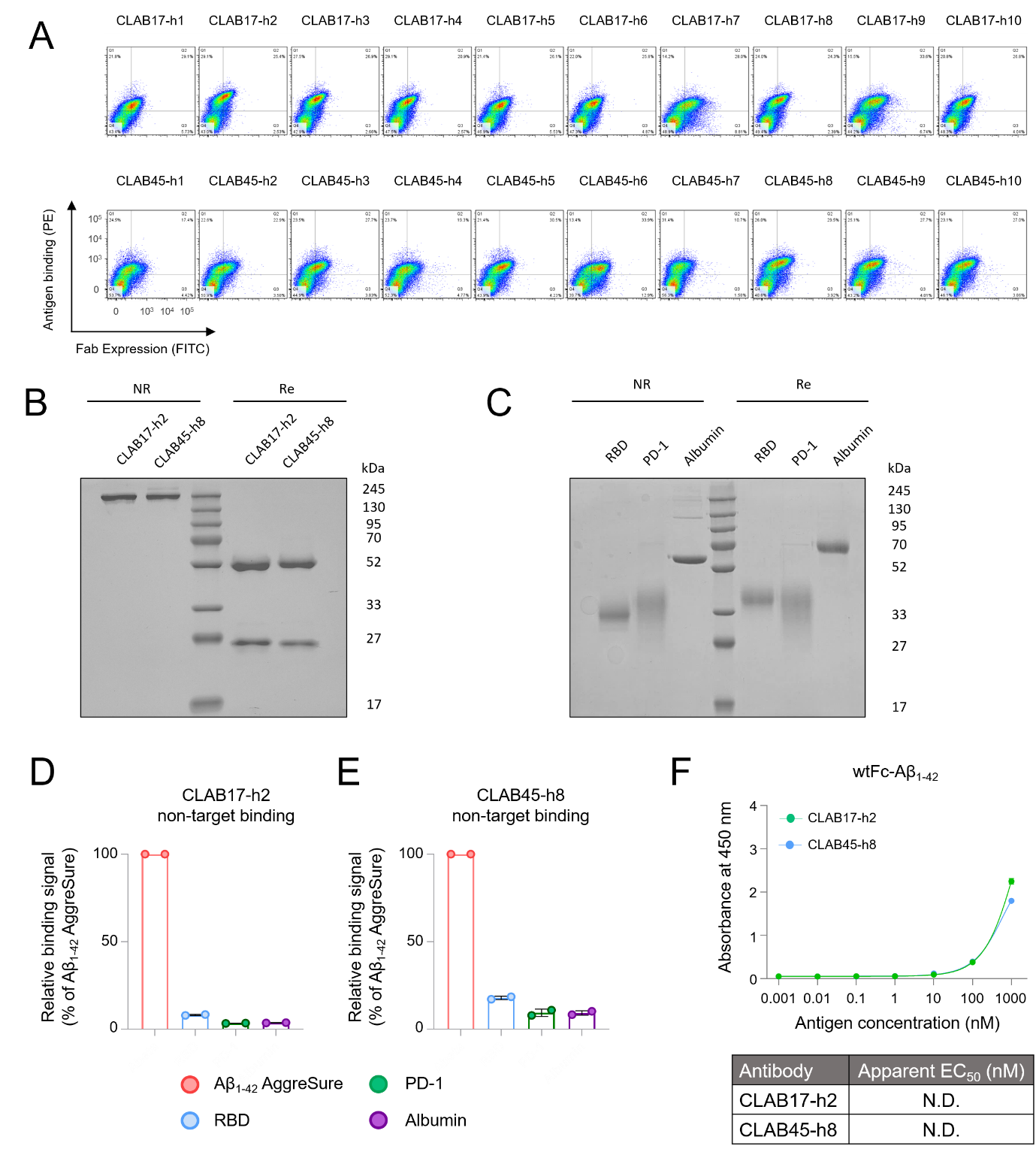

**Supplementary Fig. S3. Single-clone analysis, quality control, non-target binding, and wild-type Fc (wtFc)-Aβ_1-42_ binding of humanized CLAB antibodies.** **(A)** Single-clone fluorescence-activated cell sorting (FACS) profiles of all 20 humanized variants (10 per lineage) isolated from the CLAB17 and CLAB45 focused humanization libraries, analyzed for fragment antigen-binding (Fab) expression and binding to 30 nM Aβ_1-42_ AggreSure. CLAB17-h2 and CLAB45-h8 were selected for immunoglobulin G (IgG) conversion. **(B)** Reducing (Re) and non-reducing (NR) sodium dodecyl sulfate-polyacrylamide gel electrophoresis (SDS-PAGE) of purified IgG-formatted CLAB17-h2 and CLAB45-h8, with purity estimated by ImageJ densitometry of Coomassie-stained bands; markers in kDa. **(C)** SDS-PAGE of the biotinylated non-target proteins used in (D,E): SARS-CoV-2 spike receptor-binding domain (RBD; Wuhan-Hu-1), human programmed cell death protein 1 (PD-1), and human albumin. **(D,E)** Antibody-coated enzyme-linked immunosorbent assay (ELISA) of CLAB17-h2 and CLAB45-h8 against biotinylated Aβ_1-42_ AggreSure (target-positive control) and the non-target panel, all at 100 nM. Raw optical density at 450 nm (OD_450_) was normalized to the Aβ_1-42_ AggreSure signal for each antibody without background subtraction (AggreSure set to 100%); normalized values are reported in the Results. Because the assay lacked background subtraction, irrelevant IgG controls, and matched biotin-labeling ratios across proteins, these are exploratory non-target binding controls, not a quantitative polyspecificity assessment. **(F)** Antibody-coated ELISA of CLAB17-h2 and CLAB45-h8 against serially diluted biotinylated wtFc-Aβ_1-42_ (0.001–1000 nM). Binding was right-shifted and incompletely saturating, so apparent half-maximal effective concentration (EC_50_) values were not determined (N.D.); this does not indicate absence of wtFc-Aβ_1-42_ binding. Data are mean ± standard deviation (SD).

**Supplementary Table S1.** **Primers for murine heavy-chain variable (VH) and light-chain variable (VL) amplification.** Forward and reverse primers used to amplify murine heavy-chain and kappa light-chain variable-region repertoires from immune-derived complementary DNA (cDNA).

| VH PCR | |
| --- | --- |
| Forward (5’ to 3’) | Reverse (5’ to 3’) |
| GAGGCCGCTAGGGCCGAGGTTCDSCTGCAACAGTY | CGAGGAGACGGTGACMGTGG |
| GAGGCCGCTAGGGCCCAGGTGCAAMTGMAGSAGTC | CGCAGAGACAGTGACCAGAG |
| GAGGCCGCTAGGGCCGAVGTGMWGCTGGTGGAGTC | CGAGGAGACTGTGAGASTGG |
| GAGGCCGCTAGGGCCCAGGTTAYTCTGAAAGAGTC |  |
| GAGGCCGCTAGGGCCGAKGTGCAGCTTCAGSAGTC |  |
| GAGGCCGCTAGGGCCCAGATCCAGTTSGYGCAGTC |  |
| GAGGCCGCTAGGGCCCAGRTCCAACTGCAGCAGYC |  |
| GAGGCCGCTAGGGCCGAGGTGMAGCTASTTGAGWC |  |
| GAGGCCGCTAGGGCCGAAGTGAAGMTTGAGGAGTC |  |
| GAGGCCGCTAGGGCCGATGTGAACCTGGAAGTGTC |  |
| GAGGCCGCTAGGGCCCAGATKCAGCTTMAGGAGTC |  |
| GAGGCCGCTAGGGCCCAGGCTTATCTGCAGCAGTC |  |
| GAGGCCGCTAGGGCCCAGGTTCACCTACAACAGTC |  |
| GAGGCCGCTAGGGCCCAGGTGCAGCTTGTAGAGAC |  |
| GAGGCCGCTAGGGCCGARGTGMAGCTGKTGGAGAC |  |

| Vk PCR | |
| --- | --- |
| Forward (5’ to 3’) | Reverse (5’ to 3’) |
| GATAAAAGAGAGGCCGCTAGGGCCGACAWTGTTCTCACCCAGTC | ACAGATGGTGCAGCCACCGTGCGTTTBATTTCCAGCTTGG |
| GATAAAAGAGAGGCCGCTAGGGCCGACATCCAGATGACACAGWC | ACAGATGGTGCAGCCACCGTGCGTTTTATTTCCAATTTTG |
| GATAAAAGAGAGGCCGCTAGGGCCGATRTTGTGATGACCCAGWC |  |
| GATAAAAGAGAGGCCGCTAGGGCCGACATTSTGMTGACCCAGTC |  |
| GATAAAAGAGAGGCCGCTAGGGCCGATGTTGTGVTGACCCAAAC |  |
| GATAAAAGAGAGGCCGCTAGGGCCGACACAACTGTGACCCAGTC |  |
| GATAAAAGAGAGGCCGCTAGGGCCGAYATTKTGCTCACTCAGTC |  |
| GATAAAAGAGAGGCCGCTAGGGCCGATATTGTGATRACCCAGGM |  |
| GATAAAAGAGAGGCCGCTAGGGCCGACATTGTAATGACCCAATC |  |
| GATAAAAGAGAGGCCGCTAGGGCCGACATTGTGATGWCACAGTC |  |
| GATAAAAGAGAGGCCGCTAGGGCCGATRTCCAGATGAMCCAGTC |  |
| GATAAAAGAGAGGCCGCTAGGGCCGATGGAGAAACAACACAGGC |  |

**Supplementary Table S2. In silico developability liability scan of humanized CLAB17-h2 and CLAB45-h8.** Final humanized heavy-chain variable/light-chain variable (VH/VL) sequences were evaluated for sequence-based liabilities and structure-based developability metrics using the Therapeutic Antibody Profiler (TAP). Sequence-based liabilities were assessed by motif-based inspection of Kabat-annotated VH/VL sequences. TAP metrics were interpreted according to the five computational developability guidelines proposed by Raybould et al.

| Antibody | Extra/unpaired Cys | N-linked glycosylation sequon (N-X-S/T) | CDR-resident NG/NS motif | CDR-resident DG motif | CDR-resident Met/Trp | Total CDR length | CDR vicinity PSH | CDR vicinity PPC | CDR vicinity PNC | SFvCSP | TAP flag summary |
| --- | --- | --- | --- | --- | --- | --- | --- | --- | --- | --- | --- |
| CLAB17-h2 | Not detected | Not detected | Not detected | Not detected | Met in HCDR1 and LCDR1 | 47 | 136.7 | 0.362 | 0.000 | 12.71 | No amber or red flags |
| CLAB45-h8 | Not detected | Not detected | Not detected | Not detected | Met in HCDR1 and LCDR1; Trp in LCDR3 | 48 | 145.5 | 0.000 | 0.000 | 6.00 | No amber or red flags |

N-X-S/T, N-linked glycosylation sequon in which X is any residue except proline; NG/NS, potential asparagine deamidation-associated motifs; DG, potential aspartate isomerization-associated motif; Met/Trp, methionine or tryptophan residues located within Kabat-defined CDRs and recorded as potential oxidation-monitoring sites. PSH, patches of surface hydrophobicity; PPC, patches of positive charge; PNC, patches of negative charge; SFvCSP, structural Fv charge symmetry parameter. Sequence liabilities and TAP flags were interpreted as computational triage results, not as experimentally confirmed developability liabilities.
